# Swimming motility in the gut microbiota is diverse and increased in inflammation

**DOI:** 10.64898/2026.08.18.744924

**Authors:** Maria D. Chiotelli, Charlie Pauvert, Nicole S. Treichel, Eva-Lena Stange, Kaiyi Zhang, Aline Dupont, Agnes Seeger, Naveen Krishna Kanagaraj, Aiara Lobo Gomes, Johanna Reißing, Jonas Pes, Natalia Torrow, Tony Bruns, Nurdan Guldiken, Angela Schippers, Ana Izcue, Thomas Clavel, Marianne Grognot

## Abstract

Swimming motility has long been studied as a virulence mechanism of enteric pathogens, while the resident microbiota’s motility has only been inferred from proxies or explored through a few model species. This study presents a direct, functional analysis of gut bacterial motility in health and inflammation. Using phase-contrast microscopy and high-throughput 3D tracking, we quantified motile bacteria and characterized their swimming behaviours directly in diluted fresh gut content. In healthy mice, fewer than 3% of bacteria were motile along the digestive tract, and their swimming patterns were dominated not by the run-tumble behavior of model gut species but by diverse behaviours rich in reverses. In five mouse models with intestinal inflammation (spanning chemical, genetic, dietary and infectious etiologies) the motile fraction rose by at least 4-fold, correlating with elevated Lipocalin-2 where measured. Reverse-rich patterns remained prevalent in these inflamed conditions, with the notable exception of *Salmonella* infection. Paired metagenomics and metatranscriptomics showed enrichment of flagellar genes, while communities transferred into cecal water from inflamed mice raised their motile fraction within an hour, indicating that the rise reflects both enrichment of motile taxa and rapid modulation within existing populations. In vitro assays with human-derived isolates confirmed motility across several phyla, with variability down to strain level, and identified oxygen and viscosity as key modulators. These findings support increased motility as a hallmark of the inflamed gut and challenge established assumptions about gut bacterial motility.

**Significance Statement:** Several gut pathogens swim to survive and infect, but whether the trillions of resident bacteria also swim has only been inferred indirectly. We find that swimming is rare, under 3% of bacteria in fresh gut content. By 3D-tracking individual bacteria, we show that they swim unlike the run-tumble motility of Escherichia coli, the implicit model for gut bacterial motility since the 1970s. In five mouse models of intestinal inflammation, far more bacteria swim, from both enrichment of motile species and rapid switching within existing ones. How and when gut bacteria move, not just which are present, may matter in inflammatory diseases.

## Introduction

Bacterial swimming motility, powered by one or more flagella, enables bacteria to actively navigate their environment. Navigation abilities in turn modulates bacteria’s performance^1^ such as spreading, reaching food sources, surviving, or competing with other bacteria. In the gut, motility has long been studied as a virulence mechanism of enteric pathogens, which can provide bacteria with the ability to resist clearance by flow, facilitate penetration of mucus to access host epithelial cells, and enable travel to nutrient niches^2^. The host deploys a repertoire of innate and adaptive immune countermeasures to control this bacterial behaviour, including a Toll-like receptor system (Tlr5) dedicated to the detection of flagellins, the monomeric component of the flagella.

The relationship between bacterial motility and gut health extends beyond enteric pathogens, to the trillions of commensal bacterial cells that we host. Studies have proposed that bacterial motility is altered *in vivo* under various inflammatory conditions using surrogate markers. Cullender et al.^3^ found increased flagellin levels in faeces of mouse models with adaptive immune deficiency or impaired innate immune recognition of bacterial flagellin. Increased flagellin levels were associated with altered mucus barrier, bacterial encroachment and increased gut permeability^3,4^. Tran et al.^5^ noted elevated faecal flagellin levels in obese individuals. Several studies with diverse disease models found significantly increased genes related to motility for example in human cirrhosis with bacterial translocation^6^ or in mouse colitis^7^. These results suggest a positive correlation between motile commensal bacteria and inflammatory states.

However, the studies mentioned above are indirect. For example, flagellin measurements are a poor proxy for motility levels, as motile bacteria can display different numbers of flagella per cell^8^. Despite its putative role in inflammation, there is to our knowledge no direct assessment of (i) how many gut bacteria are motile nor (ii) which swimming behaviours are displayed by gut commensals. This gap obscures whether motility and/or specific motility patterns could confer advantages in inflamed environments. Even in vitro, the characterisation of motilities of commensal bacteria is very limited, as most efforts have focused on enteric pathogens or only a few specific model commensal species^9,10^ beyond the model organism *Escherichia coli*. It is unclear, for example, if the gut bacteria display the large diversity of swimming behaviours that has been slowly uncovered in the past 15 years in other ecosystems^11^, or if they all follow the archetypal run-tumble motility, found in model gut species such as *E. coli*^12,13^ or *Bacillus subtilis*^14,15^.

To fill these gaps, we directly interrogated the swimming motility of gut commensals and association to gut inflammation. Our approach is powered by a 3D tracking method^15^ that yields long trajectories of most motile bacteria in a dilute sample, without the need for any staining. This high-throughput, stain-free method alleviates common constraints due to microbiota diversity, the anaerobic conditions required by most commensals and the expected low occurrence of motile bacteria in healthy states. We first present a functional, direct observation of motility levels and motility types, in fresh gut contents of both healthy mice and five mouse models of inflammation, as well as in two human samples. We asked whether the fraction of motile gut bacteria increases in inflammatory conditions, and whether such changes reflect altered community composition, modulation of motility expression, or both. We developed an assay under anaerobic conditions to detect the presence of motility across 60 isolates from the Human intestinal Bacteria Collection (HiBC^16^). We then systematically tested environmental modulators (e.g. blood, viscosity, oxygen) in 9 motile isolates. Overall, we provided the first direct and quantitative portrait of bacterial swimming motility in the mammalian gut, and highlight its association with inflamed gut states.

## Results

### Gut bacteria display diverse swimming behaviours

We established a method to acquire movies of bacteria from fresh gut content diluted into anaerobic PBS (Figure 1a), maintaining stable bacterial motility for nearly 11 hours after collection (SI Figure 2a,c). From the acquired movies, the fraction of motile bacteria (i.e. the percentage of bacteria swimming relative to the whole population) is estimated (see SI Discussion 1) and 3D tracks are obtained using a stain-free, high-throughput method^15^. We first applied this approach to gut content sampled along the digestive tract of 3 healthy mice (Figure 1b). The motile fraction estimates varied depending on the gut compartment, but always stayed under 3 %, meaning that less than 3 percent of the micron-sized particles observed were deemed actively motile (as opposed to passively diffusing, displaying Brownian motion).

**Figure 1.**
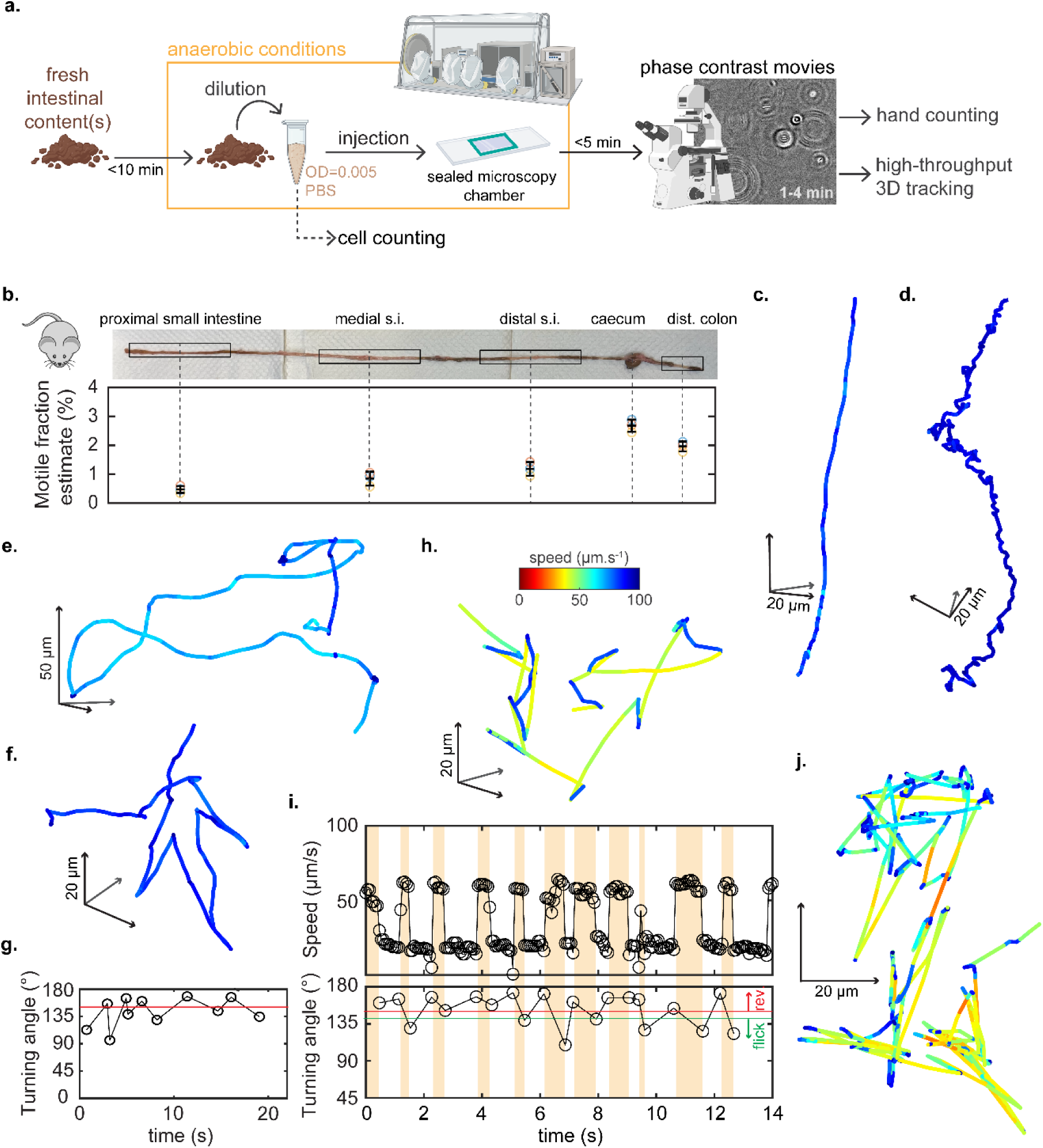
Swimming motility in gut contents from healthy mice. **(a)** Schematic overview of our protocol to estimate motile fraction and 3D track bacteria in fresh gut contents. Shortly, fresh gut content is very quickly imported into an anaerobic chamber, diluted in PBS, and injected into microscopy sample chambers that are then sealed with wax and brought for acquisition of 1-4 min movies under a phase contrast microscope. Hand-counting of motile bacteria is done to compute the motility fraction estimate (MFE), and a high-throughput 3D-tracking method is applied to extract 3D trajectories of individual bacteria (in a tracking volume of approximately 350×400×200 µm^3^). More details can be found in Methods. **(b)** Motile fraction estimates along the digestive tract of healthy mice (n=3). The intestines were taken, imported into an anaerobic chamber then cut in 5 regions of interest (picture above), and treated as shown in panel a to compute the motile fraction estimates (panel below) and 3D track bacteria. **(c)** Bacterial trajectory without turn. **(d)** Bacterial trajectory with cell wobbling. **(e)** Run-tumble bacterial trajectory. **(f,g)** Bacterial trajectory, with turn analysis showing an alternance of turning angle suggesting a run-reverse-flick motility. **(h,i)** Bacterial trajectory, with instantaneous speed and turn analysis suggesting a run-reverse-flick motility with runs of alternating speeds. (**j)** Reverse-rich bacterial trajectory. For all trajectories, color-coding represents instantaneous speed, and arrows indicate the position and scale in micrometres.

**Figure 2.**
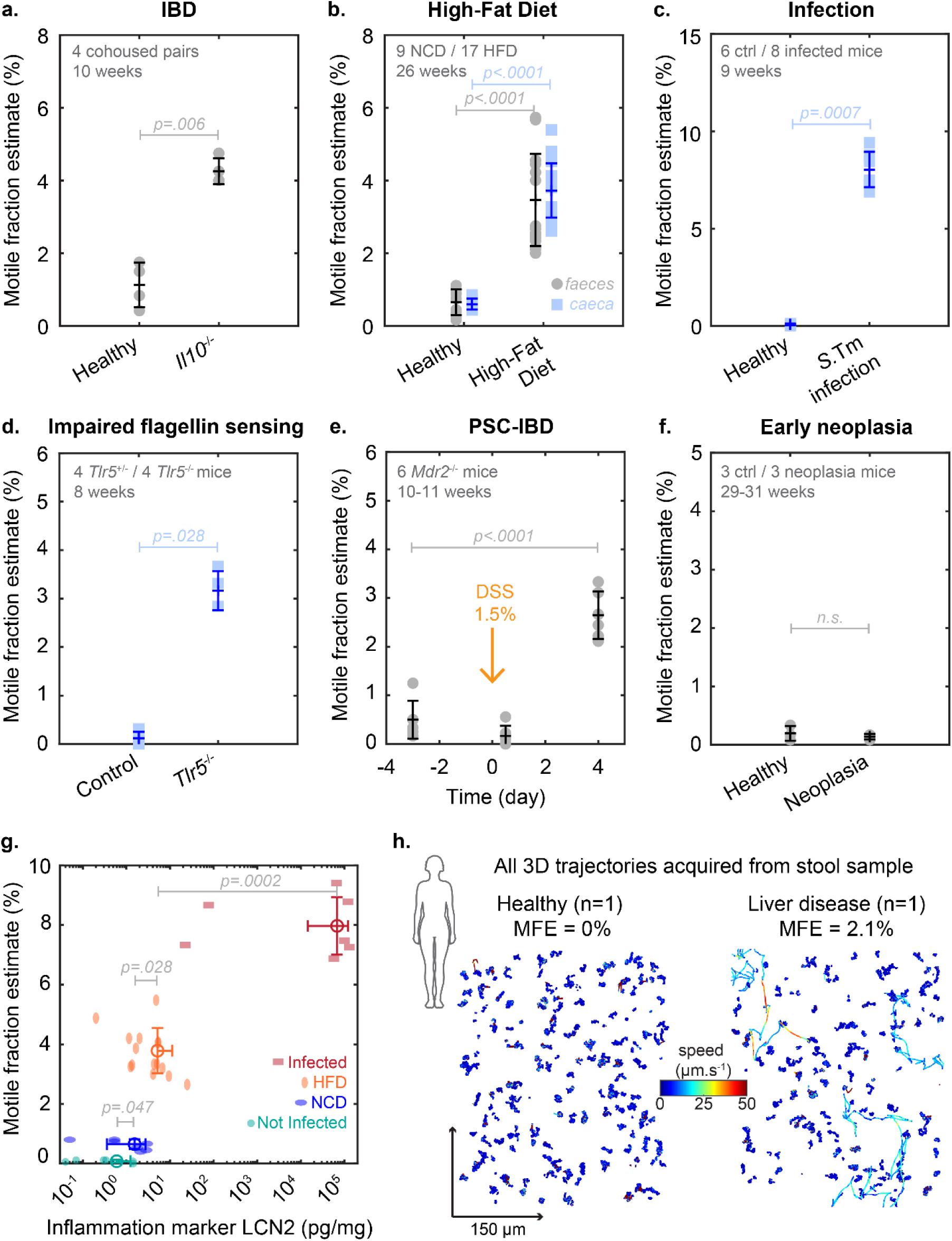
Motile fraction estimates increase in 5 mice models characterised by intestinal inflammation (IBD, PSC-IBD, High-Fat Diet, Infection, Impaired flagellin sensing) but not in a mouse model of intestinal neoplasia without colitis. (a, IBD) MFE in faeces of cohoused pairs of control and *Il10*^−/-^ mice (n=4 pairs, aged 10 weeks) showing a 3.8-fold increase. **(b, HFD)** MFE in faeces and caecal content of 26-week-old mice fed either a normal chow (n=9) or high-fat diet (n=17) showing a 5.3-fold or 6.2-fold increase respectively. **(c, Infection)** MFE in caecal content of 9-week-old mice either not infected (n=6) or infected with *Salmonella enterica* serovar Typhimurium (n=8), showing a 102-fold increase. **(d, impaired flagellin sensing)** MFE in caecal content of *Tlr5*^+/-^ (n=4, control) or *Tlr*5^−/-^ (n=4) mice (8-week-old) showing a 26.7-fold increase. **(e, PSC-IBD)** MFE in faeces of n=6 *Mdr*2^−/-^ mice (10 to 11 week old) before and after introduction of 1.5% DSS in their drinking water to induce colitis, showing a 5.3-fold increase between day -3 and a day +4. (f, Neoplasia) *VilCreERT2* β*-cat*^Δ*Ex*3^*^/+^* mice either not induced (n=3) or induced (n=3) for neoplasia. Across panels a-f, each point represents a different mouse, averaging each at least 3 technical replicates. Across panels a-f, statistical comparisons were performed using a two-tailed Mann–Whitney U test (Wilcoxon rank-sum test) except for the co-housed pairs in the IBD model and the mice before/after DSS in the PSC-IBD model, compared using a two-tailed paired t-test with unknown variance. **(g)** MFE given Lipocalin-2 concentration (LCN2) in caecal contents from a subset of the animals of 2 inflamed models: HFD, orange ellipses (n=14) and *S.* Tm. Infection, red squares (n=7). Their respective controls are in blue ellipses (NCD, n=7) and green circles (n=6). Each sample (i.e. point) originates from a different mouse. Statistical comparisons were performed using a two-tailed Mann–Whitney U test. **(h)** All 3D trajectories of more than 1s acquired from a fresh stool sample from a healthy human (left) or a patient with advanced liver disease (cirrhosis, right). Trajectories are color-coded based on instantaneous speed. No motile bacteria were seen in the healthy individual material, while an MFE of 2.1±0.3% was measured in the patient’s sample (inner material) over 3 technical replicates.

Despite the low occurrence of motile bacteria, the high-throughput 3D tracking approach allowed us to extract 378 motile trajectories of more than 2 seconds, revealing a wide variety of swimming behaviours.

First, in terms of swimming speed, motile bacteria were on average slow with an average speed of 19 µm/s, never exceeding 50 µm/s. Second, in terms of swimming patterns, we observed: straight runs (Figure 1c), long runs with visible cell body rotation/wobbling (Figure 1d) and run-tumble behaviours similar to e.g. *E. coli* (Figure 1e) but even more patterns characterised by multiple reverses (sharp turns of 150° or more, Figure 1f,h,j). In these patterns with reverses, we notably found run-reverse-flick behaviours with either constant (Figure 1f,g) or alternating swimming speeds (Figure 1h,i). Others bore similarities to run-reverse-pause behaviours (Figure 1j) but many could not be easily classified in terms of archetypal behaviours previously described in the literature.

The abundance of reverse-rich patterns, as opposed to run-tumble motility with random turning angles typically associated with gut bacteria previously studied (*E. coli*, *Salmonella* spp., *B. subtilis*, etc.) was striking. We confirmed it quantitatively by trajectory analysis. An automated detection of turns developed previously^17^ was applied to all motile trajectories obtained from the 3 healthy mice (see Methods). 132 out of 378 trajectories (∼35%, against 42% by visual inspection) had at least one turn identified as a reverse, defined here as a turn by 160° or more. Overall, the large portion of trajectories with reverses suggests an at least at least as large a proportion of bacteria being motile by means of a polar flagellum, a polar tuft of flagella, or a polar flagellum at each pole^11^.

### An increased fraction of gut bacteria is motile in inflamed gut conditions

We next measured the fraction of motile bacteria in perturbed gut states, by collecting fresh faeces and/or caeca in multiple mouse models (see Methods). Five of these models are associated with gut inflammation of varying levels: the *Il10*^−/-^ model of Inflammatory Bowel Disease (IBD), the dextran sulphate sodium model of colitis in *Mdr2*^−/-^ mice mimicking Primary Sclerosing Cholangitis with IBD (PSC-IBD), the infectious enteritis model of *Salmonella enterica subsp. enterica* serovar Typhimurium (*S.* Tm infection), the choline-deficient High-Fat Diet (HFD) model of steatotic liver disease with intestinal mucosal damage, and the flagellin sensing deficiency (*Tlr5*^−/-^) model of dysbiosis with spontaneous colitis. A model of early intestinal neoplasia without visible gut inflammation was also analysed.

An average increase of at least 3.8-fold in the fraction of motile bacteria was observed in the five mouse models of gut inflammation compared to healthy controls (Figure 2a-e) but not in the intestinal neoplasia model (Figure 2f). The largest motile fraction estimate (MFE) was observed in the caecal contents during enteric infection, with roughly 16% of bacteria swimming. We then asked if the differences in MFE correlated with the level of inflammation across two models. To do so, we quantified the inflammation marker Lipocalin-2^5,18^ (LCN-2) in caecal content of two models yielding different MFE and different magnitudes in MFE changes: the diet model and the infection model. In both models, conditions with significantly increased MFE also showed significantly elevated Lipocalin-2 levels (Figure 2g), consistent with the association between motility and inflammation. Finally, we obtained fresh faecal samples from one healthy adult and one adult with liver cirrhosis and did observe more motile bacteria in the latter (Figure 2h). Sample collection did not allow for quick import into an anaerobic chamber (see Methods) thereby likely substantially underestimating motility levels (SI Figure 2c) but still providing a qualitative extension of our results to one human context of inflammation.

### Increase in motile fraction of gut bacteria can be linked to both motile taxa enrichment and rapid environmental modulation of motility within taxa

To disentangle changes in microbial community composition from changes in gene expression, we performed metagenomic and metatranscriptomic profiling in the PSC-IBD model, before (day -3) and after (day 4) introduction of DSS to induce colitis (Figure 3a), with deep, quality-filtered sequencing of both nucleic acid fractions in every sample (see Methods). The 5.3-fold increase in MFE was associated with a 2-fold increase in the presence of 21 most prevalent flagellar genes, while their transcription was heterogenous across mice, as shown in the heatmaps for differentially abundant (Fig. 3a, left) or differentially expressed (Fig. 3a, right) genes. This resulted in the motility function abundance within the entire ecosystem being significantly increased, while the increased function expression did not reach statistical significance (Figure 3b). This indicates that the quantitative increase in motility measured by microscopy was partially due to an enrichment of motile taxa. Microscopy showed a clear motility increase in every mouse, yet flagellar transcription did not change significantly and varied widely between animals. This dissociation reinforces the premise of this work: sequencing-based proxies report the genetic potential for motility, and only imperfectly its realisation. Transient, pulsed flagellar expression and taxon-specific response kinetics could both weaken the correspondence at a single timepoint.

**Figure 3.**
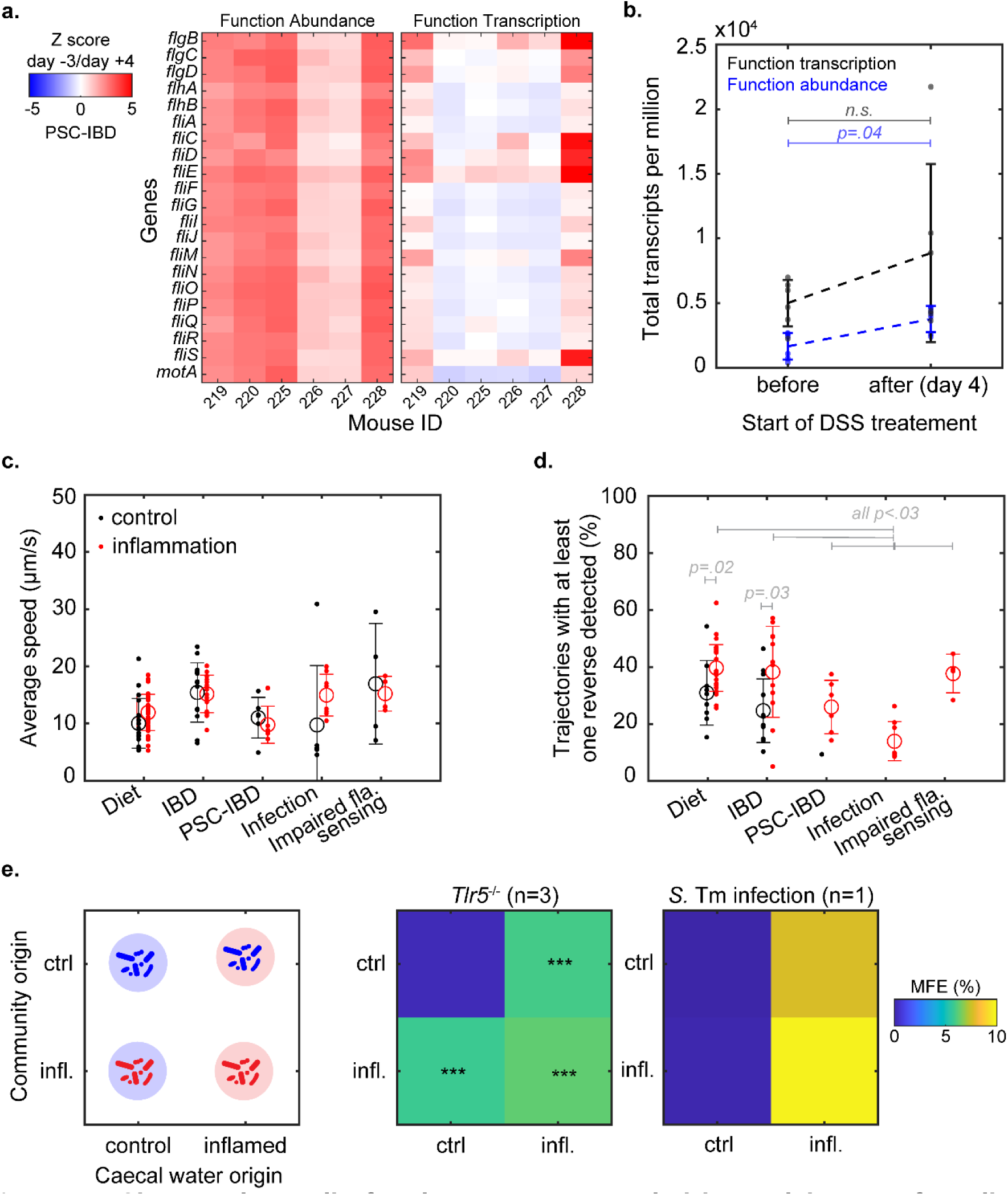
Changes in motile fraction are accompanied by enrichment of motile taxa and can be driven, within an hour, by the chemical environment. **(a)** Standardized changes (z-scores) of function abundance and function expression of 21 selected flagellar motility genes in faecal samples of PSC-IBD model, before (day -3) and after (day 4) induction of colitis by DSS treatment. Z-scores indicate by how many standard deviations each gene’s abundance/transcription deviates from its mean across all six mice pre-DSS. Genes are selected from the flagellar assembly pathway (KEGG ko02040) following a selection criterion detailed in Methods and based on later phenotyping results. **(b)** Total DNA (function abundance) or RNA (function transcription) transcripts per million summed over the 21 selected genes from the flagellar assembly pathway (see Methods), before and 4 days after addition of DSS to induce colitis in the PSC-IBD mouse model. Statistical comparisons were performed using a one-tailed Mann–Whitney U test (Wilcoxon rank-sum test). **(c)** Average bacterial swimming speed measured in each control (black dot) or inflamed (red dot) mouse, for the 5 models yielding significant increase in MFE. Averages and standard deviations are displayed as dots and error bars. **(d)** Percentage of trajectories with at least one reverse detected in each control (black dot) or inflamed (red dot) mouse, when there were at least 20 motile trajectories per mouse to analyse. Averages and standard deviations are displayed as dots and error bars. P-values originate from one-tailed Mann–Whitney U tests. **(e)** Swap experiment concept (left): from fresh caecal samples, complex communities (from inflamed or control mice) are diluted in fresh caecal water prepared from caecal content of either inflamed or control mice. Motile fraction estimates (MFE) are measured between 30 to 50 min after dilution in caecal water. Samples from 3 mice (*Tlr5*^−/-^, with 3 control *Tlr5*^+/-^ mice) or 1 mouse (*S.* Tm infected, with 1 control uninfected mouse) were used. *** indicates a p-value under 0.001 relative to the control communities in control caecal water, using a paired two-tailed t-test.

To assess potential ecological factors driving motility, we asked whether motility levels can shift faster than community composition itself. To do so, we tested if the motility of bacteria from fresh caecal samples was altered within 30-50 min of contact with sterile-filtered caecal contents (caecal water) from inflamed or control mice. We did so in two models: the flagellin-detection deficient mouse model (*Tlr5*^−/-^) and the *Salmonella* Typhimurium infection model (S. Tm). Figure 3e summarises the resulting community-background swaps and their impact on each community’s MFE within 30-50 min of contact. Complex communities from control conditions showed an increase in motility when put in contact with caecal water from inflamed conditions, i.e. *Tlr5*^−/-^ mice (n=3 mice) or infected mice (n=1). Complex communities from inflamed conditions showed model-dependent responses to control caecal water: motility decreased in the infection model but was maintained in TLR5^−/-^. Such difference may reflect model-specific factors (e.g. stronger immune mediators in infected samples). These results illustrate that the same bacterial community can, in under one hour, change its motility levels depending on the gut’s chemical environment.

### Run-tumble motility is not a representative motility behaviour of motile gut bacteria

We then interrogated the 3D trajectories from the complex communities in all models, to see if the motility properties differed from our previous observations in healthy mice. While the bacterial swimming speeds did not differ between inflamed conditions and their respective controls (Figure 3c), the pattern analysis showed significant enrichment of trajectories with reverse(s) in at least 2 models (Diet and IBD, Figure 3d). The PSC-IBD, Infection and Flagellin sensing deficient models did not yield enough motile trajectories in their respective controls to directly compare the enrichment (i.e. less than 20 trajectories of more than 2 s per mouse). All models had between 20-40% of motile trajectories with at least one reverse, with the notable exception of the *S.* Tm infection. In the latter, the number of trajectories with reverse(s) dropped significantly compared to all other inflamed conditions (Figure 3d), with a clear representation of run-tumble motility. Reverse-rich trajectories were also observed in the human stool sample from a patient with advanced liver disease (Figure 2h). These results, outside of the inflammation due to *S.* Tm. infection, highlight the relevance (through their strong representation) of reverse-rich patterns not only in healthy conditions but also in inflamed conditions.

### Phylogenetically distant gut bacteria are motile and show variability down to strain level

We then turned to in vitro work with single-isolate cultures from a human-derived collection (HiBC^16^), using two complementary approaches: a phenotypic assay to establish whether an isolate is capable of motility at all, and 3D tracking to quantify the fraction of motile cells within a population, i.e. to identify what elicits/represses motility in isolates that carry the genes but do not always express them.

The 340 isolates of this collection have been fully sequenced, allowing us to retrieve for each isolate the number of genes related to flagellar motility (see Methods). This gene screening resulted in two clear clusters: isolates carried either 0–7 or 24–42 unique genes of the KEGG pathway 02040 (“Flagellar Assembly”), with none in between. In parallel, we created a simple phenotypic assay, screening the ability of bacteria to spread through soft agar hydrogels using flagellar motility (Figure 4a, Methods), adapted from the soft agar swim plate^19–21^ to the diverse and mostly-strictly-anaerobic gut bacteria, so as to detect their ability to swim even in a small sub-fraction of the population.

**Figure 4.**
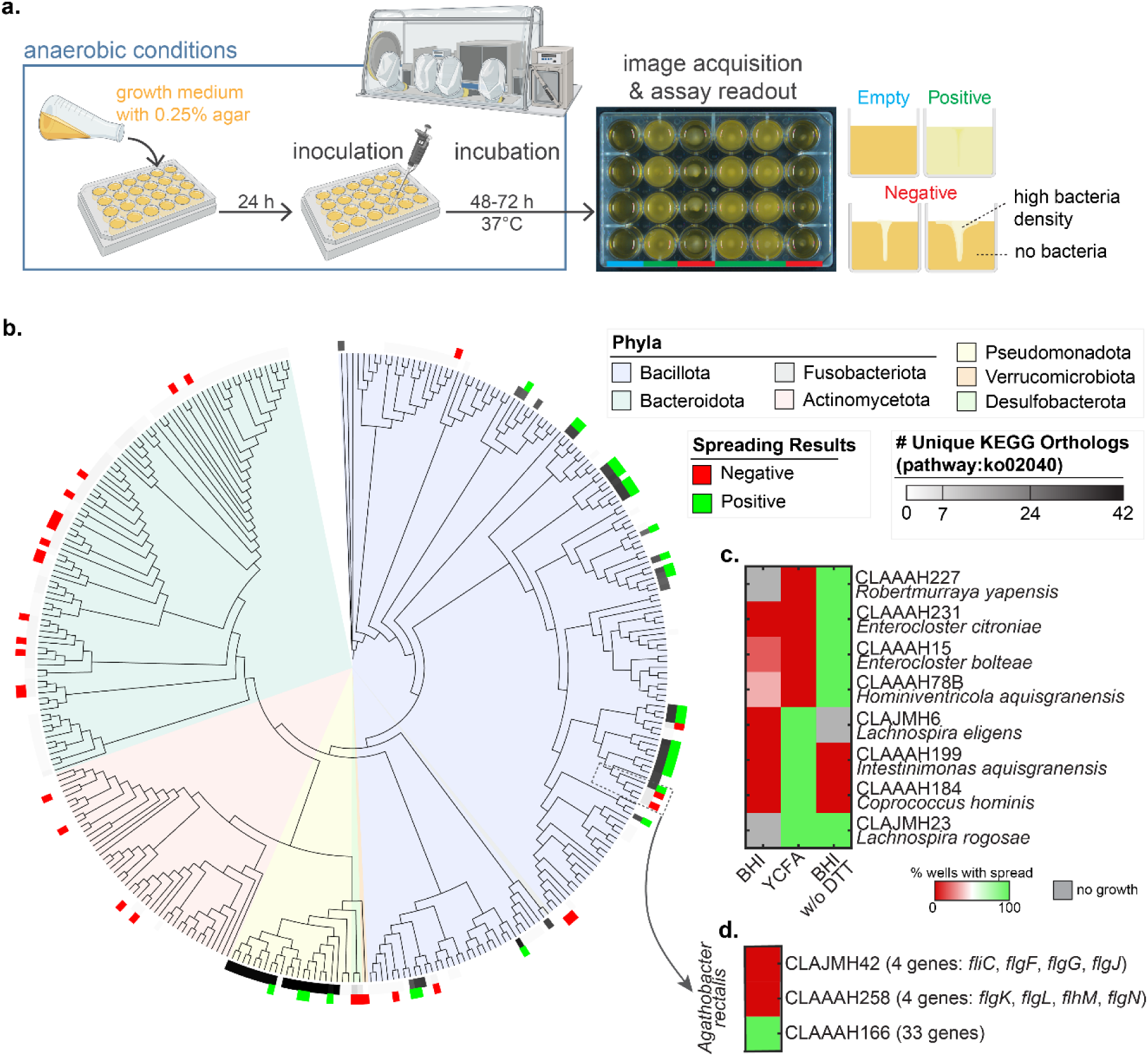
Swimming motility screening in the Human intestinal Bacterial Collection (HiBC^16^). **(a)** Schematic overview of the Simplified Soft-Agar Plate Assay (SSAPA) developed to phenotypically screen HiBC in anaerobic conditions. Growth medium (BHI, YCFA or BHI without DTT) with 0.25% agar is autoclaved and imported into an anaerobic chamber, then poured into a 24-well plate. A small volume of saturated bacterial culture(s) is injected in the centre of each well, typically 4 wells per isolate. After incubation at 37°C for 48-72h, a top-view image of the plate is taken; wells appearing bright yellow/opaque signal that the isolate can swim through the agar (positive result, green), as opposed to wells with opacity limited to growth at the injection site (negative result, red). An isolate is considered motile if any clear spread is observed in any well of any three growth conditions (BHI, YCFA or BHI without DTT). Panel partially created with BioRender.com. **(b)** Phylogenetic tree of the full HiBC collection (340 isolates) of which 60 were phenotyped: spreading results are displayed in green or red, respectively considered positive and negative for motility. This phenotypic screening matched in every case the gene screening predictions based on detection of gene orthologs from the KEGG pathway “flagellar assembly” (see Methods). **(c)** Spreading results of the 8 motile isolates showing spread in agar with growth media YCFA and/or BHI without DTT, when agar with BHI was not eliciting reliable spreading (7 isolates) or even growth (1 isolate, CLAJMH23). These 3 media were therefore sufficient to reach 100% true positive detection with the SSAPA. No false negative was detected. **(d)** Spreading results of 3 *Agathobacter rectalis* isolates along with the unique gene orthologs of KEGG pathway 02040 present in its non-motile forms.

We phenotyped 60 isolates, 29 and 31 from each cluster respectively (see Methods). Prediction and phenotype matched in both directions (Figure 4b): every isolate with at least 24 flagellar assembly genes spread in at least one of three growth conditions, and no isolate with fewer than 8 genes ever spread in any of the five conditions tested (SI Figure 4). We therefore considered motility prediction from gene screening as ground truth for presence or absence of swimming motility capabilities in further analysis, and provide in SI Figure 3 the details of the link between gene orthologs of the KEGG pathway and actual presence of motility. Detection nevertheless depended strongly on the medium. Anaerobic BHI, a standard rich medium to grow anaerobes, was taken as basal medium, but 8 of the 31 motile strains only spread in BHI without the strong reducing agent DTT or in YCFA (Figure 4c). This complete absence of spreading in BHI, lifted by other media, hints at a very tightly regulated expression of motility.

Motility was represented across several phyla, but with a high variability down to strain level. Motility had a total occurrence of 16% over the 340 isolates of the collection. The most motile phylum was the Pseudomonadota (ex-Proteobacteria) with 83% motile isolates, followed by 21% in the Bacillota (ex-Firmicutes). No isolate from the Bacteroidota or Actinomycetota phyla was motile despite a large representation in the collection. We observed variability in presence or absence of motility within genera (e.g. *Klebsiella*, *Roseburia*, *Coprococcus*) and down to strain level (i.e. between different isolates of the same species) that we phenotypically confirmed in three *Agathobacter rectalis* (Figure 4d).

### Rheology and oxygen are major effectors of swimming motility

We further explored the motility of gut commensals in vitro, as well as its modulation by chemical or rheological conditions. To do so, we selected 8 isolates from the HiBC, chosen to represent diverse species across the *Pseudomonadota* and *Bacillota* phyla. We included an additional isolate of *Proteus mirabilis*, which we isolated from a stool sample of a mouse in the HFD model (see Methods). The isolates were grown in different conditions up to an OD_600nm_=0.5-0.9 then 3D tracked, to compute the motile fractions (Fig. 5a, left). In anaerobic growth medium BHI most isolates had motile fractions under 10%. Out of all in-vitro changes in growth conditions tested, presence of oxygen (9-12%) or presence of the viscous polymer PVP K90 (4%, approx. 24 cP) yielded the highest number of strains with more than 5-fold more motile bacteria relative to the BHI condition (Fig. 5a, right). Similarly, out of the 7 motile strains able to grow but not to spread in soft agar with anaerobic BHI, 5 strains spread when traces of oxygen (estimated a few tenth of percent) were present (Figure 5c). A similar trigger of spreading upon addition of oxygen traces when grown in YCFA instead of BHI was observed in 4 out of these 5 isolates.

**Figure 5.**
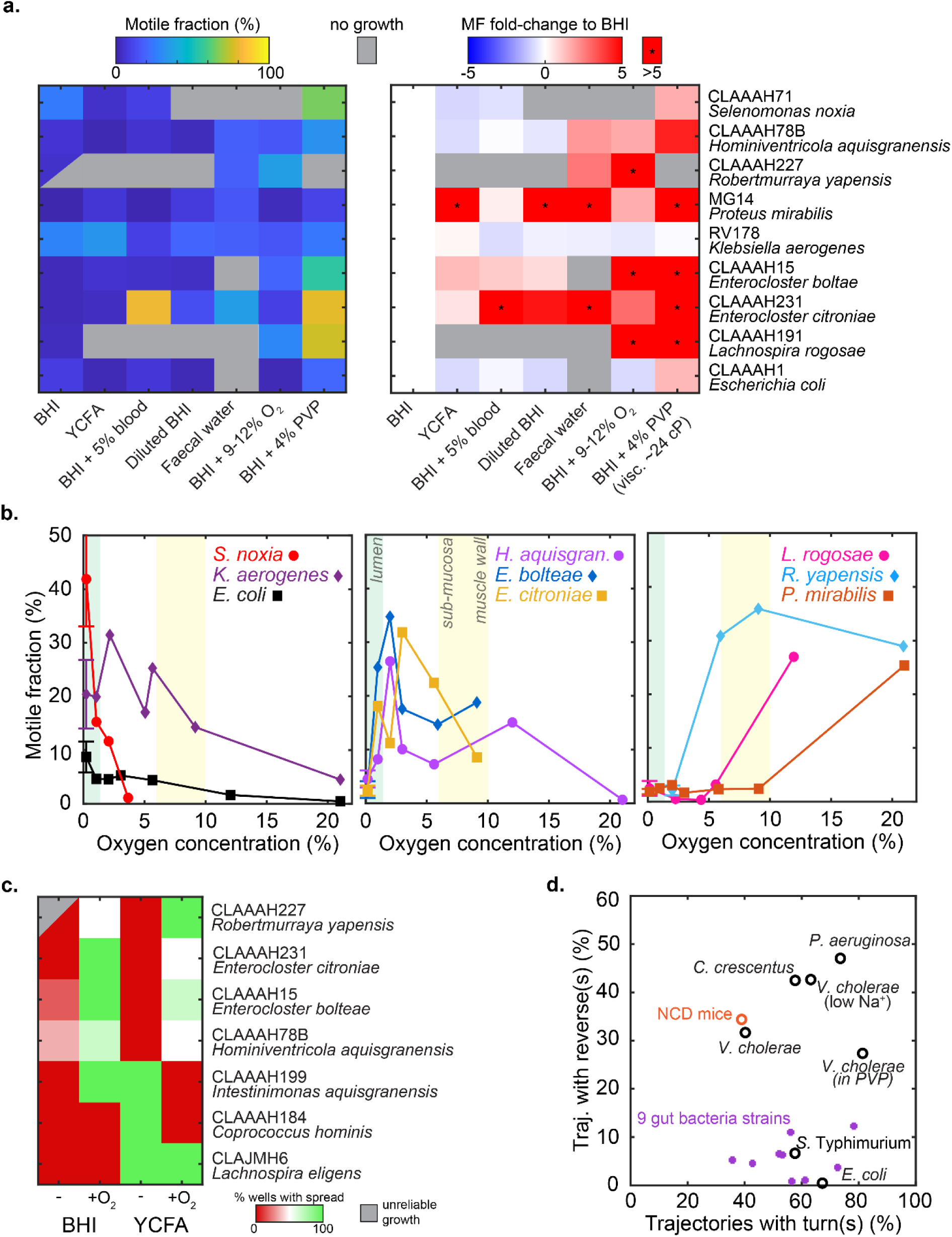
Oxygen and viscosity as major modulators of swimming motilities of gut bacteria isolates. **(a, left)** Motile fraction after growth in 6 anaerobic conditions (BHI, YCFA, BHI + 5% blood, BHI diluted by 10 in ultrapure water, BHI with viscous agent PVP K-90 at 4%, and faecal water prepared from HFD mice samples) and one with 9-12% oxygen in BHI. **(a, right)** Changes in motility levels relative to anaerobic BHI growth conditions. Conditions with more than 5 times more motile bacteria than in anaerobic BHI are signalled by a * symbol. *R. yapensis* did not always grow in pure anaerobic BHI conditions but we kept the data when it did to ensure normalization. **(b)** Motility fraction as a function of oxygen content during growth in BHI medium in nine isolates. Experiments always included a control in anaerobic conditions (<0.2% O_2_) except for *R. yapensis* (light blue diamond symbol, 2% O_2_, due to unreliable growth in anaerobic conditions). Controls were averaged over at least 3 independent experiments per strain with error bars indicating the standard error of the mean. **(c)** Spreading results of 7 motile isolates showing no spread in anaerobic BHI agar, when oxygen traces are added to BHI or to YCFA. The first 4 isolates display a clear spread only when oxygen traces (estimated at 0.2-0.4%) are present, in both YCFA and BHI. **(d)** Fraction of trajectories with at least one reverse against the total percentage of trajectories with at least one turn detected. Data for species of known motility patterns were analysed, and are displayed in black: *Caulobacter crescentus* (public dataset from Ref. ^35^, run-reverse-flick motility), *Pseudomonas aeruginosa* (run-reverse-pause), *Vibrio cholerae* in buffer or in viscous PVP K90 solution (public dataset from Ref. ^17^, run-reverse-flick motility), *Vibrio cholerae* at low Na^+^ concentrations strongly reducing its speed and flick probability (public dataset from Ref. ^17^, run-reverse motility), *Salmonella Typhimurium* (run-tumble motility) and *Escherichia coli* (run-tumble motility). Data for 9 isolates tested in panels a are displayed in purple. The reverse detection was applied for each strain to the dataset from the growth condition test (panel a) that yielded their highest motile fractions (i.e. BHI+4% PVP K-90 for all, except for RV178 and CLAAAH78B in YCFA and BHI+9.1% O_2_ respectively).

We explored in further detail the changes in motility levels across physiological oxygen concentrations. 9 isolates were grown in BHI with oxygen percentages ranging from under 0.2% (classified as anaerobic, approx. ppO <1.5 mmHg) up to 21% (similar to air, ppO ≈ 160 mmHg), and tracked over the ranges where they could actively grow (Figure 5b). Motile fraction in all populations showed significant changes with increasing oxygen concentrations, distributed equally over three apparent behaviours: decreasing, increasing or presenting a local optimum. Notably, 6 out of 9 isolates reached their highest motile fraction at an oxygen concentration above anaerobic conditions. In the inflamed gut, the lumen is expected to have higher oxygen content^22^, closer to epithelial levels (physiological ranges are displayed in Figure 5b), a range where at least 3 strains display an optimum in motility levels.

Finally, we asked whether any of these isolates displayed the reverse-rich patterns so prominent in gut contents. Applying the reverse detection to the growth condition yielding the highest motile fraction for each strain (see Methods), we found that none of the 9 isolates displayed swimming patterns rich in reverses, in contrast to both the gut communities and the reverse-rich species used as references (Figure 5d), leaving the identity of the reverse-rich gut swimmers open.

## Discussion

In this study we combined a stain-free, high-throughput 3D tracking method with sequencing and isolate-based phenotyping to obtain the first direct and quantitative portrait of bacterial swimming motility in the mammalian gut. Across five mouse models of intestinal inflammation we observed at least a 3.8-fold and up to 102-fold increase in the motile-fraction estimate (MFE) relative to healthy controls. The elevated MFE was accompanied by elevated Lipocalin-2 in the two models (four mice groups) where it was measured. While this work cannot claim that bacterial motility causes intestinal inflammation, their consistent association across multiple models suggest a contributory role that warrants future mechanistic investigation. Another key limitation is that motility was measured in diluted samples, not in native gut conditions. The caecal water swap experiments showed that dilution of communities in their own caecal waters yielded results similar to PBS (Figure 3e), and MFE was stable in PBS from 20 min on (SI Figure 2c). But we cannot exclude the possibility that dilution from dense and rheologically complex media alters motile fractions within minutes relative to in situ values. When technically accessible, future studies using intravital imaging or undiluted gut contents would significantly validate and extend these findings.

Our 3D trajectory analysis revealed a surprisingly rich repertoire of swimming behaviours in gut contents. While run-tumble (the canonical pattern of the model organism *Escherichia coli*, *Salmonella* spp. and *Bacillus subtilis*) was present, it accounted for a minority of trajectories. Instead, run-reverse, run-reverse-flick, run-reverse-pause and many other unidentified patterns rich in reverses, dominated both healthy and most inflamed conditions (≈ 30–40 % of trajectories). This calls into question the association commonly drawn between reverse-rich patterns (esp. run-reverse-flick) and a marine lifestyle^23^. Reverse-rich swimming typically arises from a low number of polar flagella rather than peritrichous flagellation, which may itself be advantageous in the gut. A reduced number of flagella could reduce the exposed flagellins compared to the peritrichous flagella, a likely advantage in the gut under close immune surveillance. Might their prevalence also point to advantages they bring for swimming through the lumen and the mucus? We believe our work demonstrates a critical need to expand motility studies beyond the *E. coli* model, and to develop quantitative assays that can capture reverse-rich motilities’ dynamics in environments that more faithfully recapitulate the gut conditions (e.g. bacteria-dense mucus with an oxygen gradient).

The absence of reverse-rich motility in the 9 bacterial isolates we screened means we cannot yet propose a new model of gut bacterial motility for now. A few gut bacteria taxa that we know can exhibit run-reverse or run-reverse-flick behaviours are members of the *Desulfovibrio*, *Helicobacter* and *Campylobacter* genera. Another intriguing possibility is that species known to display numerous flagella and run-tumble motility in the lab are not phenotypically representative of strains/isolates from the same species found in the host environment. Notably, a run-reverse-like behaviour has been reported in one of the original *E. coli* K-12 clones^24^. Another hypothesis could be that the chemical and mechanical conditions in the gut can alter the motility pattern of the same isolate compared to in-vitro. We even tested whether shear stress (an important mechanical component of the gut dynamics) could push a peritrichous species (S. Tm) towards reverse-rich patterns, but found no such effect (SI Figure 5).

Our work also highlights that the observed rise in motile fraction is likely driven by two cumulative mechanisms: (i) the enrichment of taxa with the ability to swim and (ii) transient, environment-dependent increase in motility of the same bacteria. The rapid increase in MFE observed when naïve communities were incubated with caecal water from inflamed mice supports the idea that environmental cues can quickly modulate flagellar activity in the gut. A response on a 30–50 min timescale falls at the low end of current estimates of the time required to build a flagellum de novo in order to support motility in *Salmonella* Typhimurium^25^. In vitro experiments with a panel of gut isolates revealed that oxygen concentration is a major modulator of swimming motility, significantly expanding a few earlier reports of similar effect in single-species tests^10,26^. The inflamed gut is known to become more oxygenated due to compromised epithelial barrier function and increased epithelial oxygen leakage^22,27^. Thus, we hypothesise that elevated luminal oxygen in inflamed conditions can in part drive early increase in bacterial motility. As motility fraction seemed to be very tightly regulated across gut bacteria, we envision that future large-scale screenings of motility levels may be a window into the ecological and disease contexts in which movement confers a fitness advantage.

Overall, our work invites microbiologists, (bio)physicists and immunologists to collaboratively dissect how bacterial motility shapes and is shaped by the gut ecosystem, ultimately guiding the development of interventions that tame inflammation by modulating bacterial behaviour, rather than phylogeny or abundance alone.

## Materials and Methods

### Bacterial strains, media and anaerobic cultivation

Gut bacteria strains were sourced from the HiBC, a substantial and fully sequenced collection of human intestinal bacteria (see SI Table 1). To revive the anaerobic collection, 150–200 µL of cryostock was inoculated into 9 mL Hungate tubes containing either Brain Heart Infusion (BHI), Yeast Casitone Fatty Acid (YCFA), Wilkins–Chalgren Anaerobe (WCA), or modified Gifu Anaerobic Medium (mGAM), whichever preferred medium was indicated by the HiBC repository. All exact recipes can be found in References^16,28^. Inoculation was performed using a sterile syringe and needle inserted through the rubber stoppers, which were disinfected by flaming following the application of ethanol. The aerophilic strain *R. yapensis* was revived by plating cryostock onto LB Lennox agar and incubating under aerobic conditions. A single colony was subsequently selected using a sterile loop and used to inoculate 3 mL of LB Lennox broth. All revival cultures were incubated in anaerobic conditions at 37 °C for a minimum of 18 h and, depending on growth rates, maintained for up to 96 h to ensure sufficient biomass. Revival cultures could then be used for further culturing steps depending on the experiment. Caecal and faecal waters were prepared as follows. Approximately 1 g of stool or scrapped caecal content was mixed with 4 mL of ultrapure water, thoroughly homogenized, then centrifuged at 12,000 g for 3 minutes to pellet the debris. The supernatant was sterile filtered through a 0.2-µm filter. In the few aerobic experiments, other media were used: TB (Terrific Broth; 1% Bacto Tryptone, 0.5% NaCl, pH 7) and TG (TB with 0.5% (wt/vol) glycerol)

### Lipocalin-2 ELISA

Caecal contents from the *S.* Tm infection model were collected and frozen at -80°C until further use. Caecal contents were weighed and resuspended in 1 mL PBS/0.01% Tween by vortexing for 10-15 min at room temperature^18^. The suspension was centrifuged for 10 min at 12,000g at 4°C to pellet debris and the supernatants were transferred into new tubes. Lipocalin was measured by DuoSet ELISA (Biotechne/R&D Systems, DY1857-05) according to the manufacturer’s instructions; samples were diluted 1:2, 1:5 or 1:10. Samples were read with a Spectra Max Microplate reader and lipocalin content of the homogenates were calculated using a 4-PPL-standard curve generated by the device’s software. Lipocalin levels were normalised to the weight of the used caecal content.

### Gene Screening of the Human intestinal bacteria collection (HiBC)

We annotated the 340 high-quality genomes of HiBC^16^ with bakta v1.9.4^29^ using the full database v5.1. Bakta-generated proteins sequences were screened against the genes in the motility (02040) and chemotaxis (02030) pathway from the KEGG database^30^. We used kofamscan v1.3.0 with the HMM profiles from Kofam database (2024-10-30) to detect KEGG Orthologs genes using the predefined score thresholds. Overall, the motility pathway consists of 53 genes distributed in 52 non-redundant genes, while the chemotaxis pathway consists of 16 genes distributed in 8 non-redundant genes. Screening code is available as a Snakemake^31^ workflow at https://git.rwth-aachen.de/cpauvert/motility-chemotaxis-hibc. Protein sequences of genes in plasmid sequences were obtained using the same procedure except with a parameter tuning adapted to circular sequences, however, no plasmid-borne proteins matched the motility and chemotaxis pathways in our 340 isolates. Eight genes were not detected in any isolates: K24344 (flgO), K24346 (flgQ), K24343 (flgT), K10943 (flrC, fleR, flaM), K10564 (motC), K10565 (motD), K21217 (motX) and K21218 (motY).

### Picking strains from HiBC for phenotypic validation of presence or absence of motility

The gene screening of the HiBC resulted in two clear clusters (either 0-7 or 24-42 unique flagellar assembly genes as defined by the KEGG pathway 02040). We picked 29 isolates and 31 isolates in each category respectively. Among the 29 isolates from the first category (<8 flagellar assembly genes), we elected to represent each sub-category we could observe based on which genes were found. As a result, 7 had 0 flagellar genes, 1 was the only one isolate with 7 (CLAAAH201, *D. piger*), 2 had *fliY* alone, 4 had *flgJ* alone, 1 had *flgJ*+*fliY*, 11 had *flgJ*+*motB*, and the 3 remaining were the only ones with their flagellar genes combination (including 2 displaying *fliC*). The categories and related isolates can be found in SI Figure 3. The 31 isolates from the second category (>23 flagellar assembly genes, 56 isolates total) were picked at random.

### Picking 21 genes directly associated with flagellar motility

From the gene screening of the HiBC, we extracted for each gene of the “Flagellar Assembly” KEGG pathway the frequency of its association with motility or absence of motility. The 21 genes selected are all genes found in both more than 97% of the motile isolates (i.e. isolates with more than 23 unique genes KEGG pathway 02040), and in less than 3% of the non-motile isolates (i.e. isolates with less than 8 unique genes from the KEGG pathway 02040). This ensures that we probe genes truly associated with flagellar motility, rather than genes necessary for flagellar expression but found in abundance in other processes (e.g. *flgJ*, gene for a peptidoglycan hydrolase, found in approx. 75% of motile bacteria but also in a similar share of non-motile bacteria). The 21 genes selected from the KEGG pathway 02040 were: *fliE, fliF, fliG, fliI, fliJ, fliM, fliN, fliO/fliZ, fliP, fliQ, fliR, flhA, flhB, flgN, flgC, flgD, fliC/hag, fliD, fliS, motA, fliA/whiG*.

### 24-well soft agar plate screening

Brain Heart Infusion (BHI) medium (with or without DTT) and YCFA-modified media mixed with 0.25% (w/v) agar were prepared, autoclaved, and dispensed into 24-well plates under aerobic or anaerobic conditions. After solidifying for one hour at room temperature, the plates were incubated for 24 hours at 37 °C in an anaerobic chamber (Coy Lab). Cryogenic stocks from the HiBC collection were revived, then cultures were incubated overnight in Hungate tubes at 37 °C. The following day, 0.5 µL of each revival culture was inoculated into the centre of the wells. Four technical replicates (four wells) were prepared for each strain, alongside four medium-only controls and four replicates of the negative control strain, *Shigella flexneri* (ATCC 12022). The plates were then incubated under anaerobic conditions for 48 hours. At the end of the incubation period, images were acquired using a Pixel Device (Singer Instrument Co.Ltd). A well was considered positive when: The full well turned opaque from spreading and growth and had 4 valid negative control wells on the same plate; The full well turned opaque and single-cell movement was observed under the microscope (inverted microscope Eclipse Ti2, Nikon). A well was considered negative when there was a clear sign of growth at the site of injection but no increased turbidity in the rest of the well. A well was considered inconclusive if there was no visible growth at the site of injection after 48h. Any plate where any turbidity in the empty (no bacteria) wells was noted was fully rejected from analysis.

### Motility measurements in faeces and gut content (mice)

Fresh gut content samples were collected using sterile beakers during weighing. Samples from control or inflamed conditions were collected at the same time and treated similarly. The samples were then transferred to Eppendorf tubes using sterile forceps and transferred within 10 minutes to an anaerobic chamber for processing. The quick import (under 10 min, ideally 8 min) was found to be critical for observing motility (SI Figure 2c). Caeca were opened and inverted under anaerobic conditions using sterile tweezers. Material from the inner portion of each sample was collected with a sterile loop, resuspended in sterile PBS. PBS was chosen as it yielded stable MFE over 11 hours, while communities resuspended in their own caecal water had variations within 3 hours (SI Figure 2). The optical density at 600 nm (OD_600nm_) of the solution was measured to then dilute further to a target OD_600nm_ of 0.005. Then the sample was incubated at room temperature for at least 15 min to allow adaptation to the medium, and up to 11 h. Motility chambers (∼300 µm in height), consisting of three layers of parafilm between a microscope slide and coverslip, prepared and imported in advance into the anaerobic chamber, were filled anaerobically. They were then sealed with wax (VALAP^15^) and immediately transferred outside of the anaerobic chamber to the microscope for data acquisition. For each mouse, three to five technical replicates were prepared, each originating from a different spot of the sample taken with a sterile loop, followed by an independent dilution series. Special care to alternate measurements between control and non-control conditions was taken. Each motility chamber was prepared and exported just before acquisition. Recordings were acquired on a phase-contrast microscope (see Phase contrast microscopy section). Bacterial trajectories were obtained from the acquired movies using a high-throughput 3D tracking method^15^, then filtered and analysed (see Analysis of 3D trajectories).

### Motility measurements in faeces (human)

Stool samples obtained from a healthy adult (n=1) or a patient with cirrhosis (n=1) were imported into the anaerobic chamber within 12 or 20 minutes respectively after defecation. This collection time likely lead to an underestimation of motile fraction estimates (MFE), especially in the patient’s sample. Three technical replicates (motility chambers and 2-min movies) were made after sampling the material with a sterile loop dipped into the inner part of the stool. This was crucial, as the external side of the patient’s sample yielded a lower MFE (∼1%) likely due to longer exposure to air diffusing from the outer to the inner region of the sample. Protocol was otherwise similar to that with mouse samples.

### Motility across healthy mouse gut

Gastrointestinal material was collected from five anatomical regions of three mice under normal chow diet (NCD): proximal, medial, and distal small intestine, caecum, and colon near rectum. The intestinal tract was dissected immediately after sacrifice and transferred into the anaerobic chamber for processing. Subsequent steps followed the protocol described above for motility measurements in faeces and gut content.

### Phase contrast microscopy

Motility videos were acquired at room temperature using a Nikon Ti-E inverted microscope equipped with a sCMOS camera (PCO Edge 5.5) and a 40× objective lens (Nikon CFI S Plan Fluor ELWD 40× ADM Ph2, correction collar set to 1.2 mm). Recordings were collected at 15 frames per second for 1.33-4 min (mouse models) or 2 min (in-vitro tests).

### Bacterial counting in samples

To ensure comparability across samples and experiments, bacterial concentrations were quantified in diluted suspensions used for motility acquisitions, using a haemocytometer (INCYTO C-Chip Neubauer Improved DHC-N01, NanoEntek). Briefly, a culture adjusted to an OD_600nm_ of 0.005 was mixed 1:1 with a 0.15% w/v crystal violet solution (bioMérieux), the haemocytometer chamber was loaded with the mixture, and cells were counted under a phase-contrast microscope. In case the cell concentration in samples went outside of the 1-1.5 × 10^6^ cells/ml range in one model, or showed variations within the same model, the concentration was informing a correction of the Motile Fraction Estimate (MFE) by a factor reported in SI Table 2.

### Treatment of 3D tracking data from gut contents

The 3D tracking method produced raw trajectories that recorded the frame number, spatial coordinates (x, y, z) and a goodness-of-cross-correlation metric for each frame. These trajectories were aggregated into a single structure per sample (i.e. per type of content and mouse) and smoothed using a second-order ADMM-based filter^32,33^ with a regularization parameter of λ=0.3. Each trajectory of more than 2 seconds was kept and classified as motile or non-motile using its median speed (at least 3 µm/s to be motile) then a trained model. A binomial generalized linear model (logistic regression) was fitted to a training dataset of 450 trajectories annotated as motile (150) or non-motile (300) by hand. The parameters retained as sufficient descriptors for each trajectory were: the median speed, the total start-to-end distance and the standard deviation on instantaneous speeds. The trained model was applied to 554 test trajectories, probabilities were converted into binary class labels using a 0.5 decision threshold, resulting in the confusion matrix provided in SI Figure 6a. This approach was deemed more effective for the noisier data from gut samples, compared to the typical “hard-threshold approach” used in vitro (SI Figure 6b).

### Motility of HiBC isolates in anaerobic conditions

The 9 bacterial isolates chosen from HiBC for further in-vitro tests (listed in SI Table 1) were cultured overnight at 37 °C in 9-mL Hungate tubes using BHI, BHI diluted with Ultrapure Milli-Q water (1:10), faecal water prepared from HFD mouse samples, BHI with 5% defibrinated sheep blood (Oxoid), or BHI with 4% (w/v) PVP K-90 (Sigma-Aldrich, ref. 81440). Day cultures were then inoculated at a dilution of 1:1000 from the overnight cultures into fresh 9 mL medium identical to the medium used beforehand. The cultures were grown until they reached an OD_600nm_=0.5-0.9, except in specific conditions: CLAAAH71 in YCFA (0.1), RV178 in faecal water (1), CLAAAH15 and CLAAAH231 in diluted BHI (0.46 and 0.4 respectively). The cultures were then diluted to a target OD_600nm_=0.005 in fresh medium matching there growth medium (except for BHI+ PVP, that was resuspended in BHI) and 3D tracked.

### Motility of HiBC isolates under different oxygen concentrations

We tested 9 isolates: for *S. noxia* (CLAAAH71), *K. aerogenes* (RV178), *E. coli* (CLAAAH1), *H. aquisgranensis* (CLAAAH78B), *E. bolteae* (CLAAAH15), *E. citroniae* (CLAAAH231), *L. rogosae* (CLAAAH191), *R. yapensis* (CLAAAH227), *P. mirabilis* (MG14). To establish different oxic conditions, anaerobic Hungates were opened to air for 66 seconds to 5 min without agitation, closed and homogenised. The oxygen concentrations were then measured through the septum in each Hungate, using a high-precision oxygen sensor (Microx 4, with PSt 7 probe in a syringe casing, PreSens). Day cultures were then inoculated at a dilution of 1:1000 from the overnight cultures into fresh 9 mL BHI under the same oxygen conditions. The cultures were grown until they reached an OD_600nm_ comprised between 0.70 and 0.92. For 3D tracking, the day cultures were diluted to an OD_600nm_ of 0.005 in fresh BHI with oxygen concentrations matching the growth conditions.

### Treatment of in-vitro 3D tracking data

Data was collected and organized by sample, with each sample comprising at least three technical replicates. The 3D tracking method produced raw trajectories that recorded the frame number, spatial coordinates (x, y, z) and a goodness-of-cross-correlation metric for each frame. These trajectories were aggregated into a single structure per sample and smoothed using an ADMM-based filter^32^ with a regularisation parameter of λ=0.3. Each trajectory of more than 2 seconds was kept and classified as motile or non-motile using its median speed and 2D effective diffusion coefficient (D_eff_), the latter computed using the msdanalyzer class written by J.-Y. Tinevez^34^. Shortly, D_eff_ is estimated from the mean squared displacements (MSD), using only the – coordinates to minimize noise contributions from the -axis. The MSD is fitted linearly on the first 25% of the forward trajectory and on the reversed last 25% of the trajectory, and D_eff_ derived from the slopes, with the final value taken as the mean of the two. A trajectory was classified as motile if its D_eff_ exceeded 3 µm²/s (more than 10 times above the expected D ≈ 0.2 µm²/s of a purely diffusing 1-µm particle according to the Stokes-Einstein equation) and its median speed exceeded 3 µm/s. The time-based motile fraction (total time spend swimming over total duration of all tracks) and the mean bacterial swimming speed (average of average individual speeds in the motile population) were then computed. For growth in BHI, at least 3 biological replicates in anaerobic conditions (< 0.2% O ) were aggregated into a single representative average data point per strain.

### Reverse-detection analysis in motile trajectories

The turning event detection, that we previously applied to another *V. cholerae* strain (including sharing of the code)^17^, is based on the local rate of angular change, computed from the dot product between the sums of the three consecutive velocity vectors preceding and subsequent to a time point. The threshold for a turn to begin was set here at a 5-fold rate relative to the median rate of angular change of the run segments, instead of a 7-fold rate as previously used, as visual inspection determined this low threshold to be necessary to detect reverses in the noisiest trajectories. This lower threshold leads to an over detection of low-angle turn (typ. 0-90°), as a concession to detection of almost all reverse, but these low-angle turns are rejected from the analysis. The turn ends with at least one time point under this threshold. The 3D turning angle for a turn beginning at frame *i* and ending at frame *j* is computed as the angle between the sum of the instantaneous velocity vectors at frames *i*-3 and *i*-1 and the sum of those at frames *j* and *j*+2. Reverses were defined here as turns by 160° or more. This reverse-only analysis differs from previous turn analysis as it typically over detects “turn” events at low angle (typ. 0-90° range) before rejecting them.

### Motility of model species for comparison in reverse-detection analysis

Analysed data for species of known motility patterns (with reverse-analysis displayed in Figure 5d) were mostly sourced from public datasets: *Caulobacter crescentus* (from Ref. ^35^, run-reverse-flick motility), *Vibrio cholerae* in buffer or in viscous PVP K90 solution (from Ref. ^17^, run-reverse-flick motility), *Vibrio cholerae* at low Na^+^ concentrations strongly reducing its speed and flick probability (from Ref. ^17^, run-reverse motility). We added our own datasets for *Salmonella* Typhimurium (run-tumble motility), *Escherichia coli* (run-tumble motility) and *Pseudomonas aeruginosa* (run-reverse-pause). *E. coli* and *P. aeruginosa* were plated on 2% LB agar, then a colony was picked for creating an overnight culture incubated at 30°C 250 rpm for 16h, diluted 1:200 into TG then grown at 30°C 250 rpm to a respective OD_600nm_ of 0.72 and 0.21, and finally diluted in fresh TG. *Salmonella* Typhimurium (ATCC14028) was grown directly from stock in LB up to saturation, diluted 1:200 in TB and grown up to OD_600nm_=1.05 at 30°C 250 rpm, and finally diluted in fresh MotM^35^. All strains were diluted to a target OD_600nm_=0.005 and brought to the microscope for acquisitions at 15 fps (*E. coli, S.* Tm) or 30 fps (*P. aeruginosa),* 15 to 30 minutes after dilution to allow for adaptation.

### Turn analysis for run-reverse-flick example trajectories

In Figure 1 are displayed two run-reverse-flick trajectories. The turn analysis applied was identical to the one applied in Ref. ^17^ to *Vibrio cholerae*.

### Sequencing of stool samples before and after colitis induction

Samples correspond to day -3 and day +4 after the introduction of 1.5% DSS into drinking water of 6 *Mdr2^−/-^*mice (see also Mouse models methods’ section). As resource constraints precluded sequencing all five models, the PSC-IBD model was selected for its time-resolved design, allowing paired comparisons before and after colitis induction in the same animals.

### DNA Extraction, Library preparation and short-read sequencing

DNA and RNA from mouse faecal samples were extracted using the ZymoBIOMICS DNA/RNA Miniprep Kit extraction Kit (Zymo Research) according to manufacturer’s instructions with an elution volume of 50 µL. Shotgun metagenomic DNA libraries were prepared using the NEBNext Ultra II FS DNA Library Prep Kit for Illumina (NEB) according to the manufacturer’s instructions, using 300 ng of input DNA and an automated platform (Beckman Coulter). Enzymatic shearing to approximate 250 bp was performed for 30 min. Adaptor-ligated DNA was enriched using PCR (5 cycles) and NEBNext Multiplex Oligos for Illumina (NEB) for unique dual barcoding. Size selection and clean-up of adaptor-ligated DNA were performed using AMPure beads (Beckman Coulter). Preparation of the libraries for RNA sequencing and further steps including sequencing runs were conducted at the IZKF Core Facility Genomics (UKA, RWTH Aachen University). RNA libraries were prepared in treatment randomized batches with the NEBNext UltraExpress RNA Library Prep Kit using 200 ng of RNA and including rRNA depletion for bacteria. Libraries were indexed using NEBNext Multiplex Oligos for Illumina (NEB) for unique dual barcoding. For quality assurance RNA libraries were spiked with 1 µl of a 1:250 dilution of ERCC RNA Spike-in Control Mixes (Invitrogen) each. Resulting RNA and DNA libraries were quantified (Qubit Flex, Invitrogen) and quality checked (TapeStation 4200, Agilent Technologies) before sequencing. The 24 metagenomic and metatranscriptomic libraries were each sequenced on a NovaSeq6000 (Illumina) using NovaSeq 6000 Reagents S1 v1.5 (2 × 150 cycles).

### Analysis of metagenome and metatranscriptome data

The analysis pipeline MIntO v2.4.0 was used for pre-processing and analysis of the metagenomic and metatranscriptomic reads^36^. Quality filtering and host DNA removal resulted in 56.650.337 ±30.309.485 high-quality reads for the metagenomic data, and in 28.281.779 ±7.199.388 high-quality reads for the metatranscriptomic data after host DNA and ribosomal RNA removal. The pipeline was run in genome-based mode using the reference genomes of the integrated mouse gut catalog (iMGMC^37^). Genes of the iMGMC genomes were predicted using Prokka v1.14.6. For functional annotation to the Kyoto Encyclopedia of Genes and Genomes (KEGG) kofamscan v.1.3.0 was used. Gene and transcript abundance were determined by alignment of the high-quality reads to the reference genomes. Normalization to transcripts per million (TPM) and calculation of gene and function expression profiles was conducted as described by Saenz et al. ^36^.

### Mouse models

Samples in this manuscript were all sourced from mouse experiments where they would have otherwise been discarded, without incurring changes in approved protocols. Animals were all bred locally at University Hospital RWTH Aachen. All animal experiments were conducted in accordance with institutional and national guidelines for the care and use of laboratory animals and were approved by the state licensing authorities for animal experimentation, the State Office for Consumer Protection and Nutrition of North Rhine-Westphalia (LAVE, i.e. Landesamt für Verbraucherschutz und Ernährung, North Rhine-Westphalia, Germany). Authorisation numbers and detailed information are listed in the Supporting Information.

### Human samples

Stool sample from a patient with advanced liver disease hospitalized for decompensated cirrhosis (Child-Pugh B9) or from a healthy volunteer were collected using the Gut Alive Microbiome Collection Kit (MicroViable GmbH, Germany), which was processed within 20 minutes in an anaerobic chamber. This study was conducted in accordance with the ethical principles of the Declaration of Helsinki and Good Clinical Practice (ICH-GCP) guidelines. It was approved by the Ethik-Kommission an der Medizinischen Fakultät der RWTH Aachen (EK 327/19 with protocol amendment v1.2), and written informed consent of the patient was obtained prior to sample collection.

## Acknowledgments

This work was funded by the German Research Foundation (DFG) – Project-ID 403224013 – SFB 1382. This work was partially funded by RWTH Aachen Faculty of Medicine START-Program, Großantrag awarded to N.T.

