## Supplementary Information for "Swimming motility in the gut microbiota is diverse and increased in inflammation"

**Supporting Information Text**

**SI Discussion 1: on motile percentage and motile percentage estimate.**

We define the motile fraction as the total duration of trajectory time of motile bacteria divided by the total duration of trajectory time of all bacteria tracked. This definition is used in all analysis of single-species cultures in vitro. But the increased noise in 3D tracking of diverse bacteria in gut contents, as well as the difficulty of distinguishing very slow motile bacterial tracks from non-motile tracks, limited our confidence in directly measuring the motile fraction from 3D tracking data. We therefore resorted to a motile fraction estimate (MFE), proportional to the number of motile bacteria as directly seen in the movie acquisitions, counted by hand, divided by the total duration of the movie.
In faeces, a consistent cell count of $C_{feces} \approx 1\text{-}1.5.10^6$ cells per ml was found across mice models when diluted to a target OD of 0.005 (SI Figure 1a-d). Our field of view during acquisition is approximately 400x350x300 µm = 0.042 mm$^3$ = 42.10$^{-6}$ ml, that is an expected visible number of 42x1.2≈50 bacteria. Assuming a motile trajectory duration of 0.5 min (close to the 28s measured for the 378 trajectories from 3 NCD mice) and non-motile duration equal to the acquisition duration, counting 1 motile bacteria per minute of movie is equivalent to a motile fraction of ~1%. Consequently, our motile fraction estimate (in %) in faeces is directly equal to the number of motile bacteria counted by minute of acquisition.
In caecal content, for the same dilution to a target OD=0.005, we systematically found a significantly higher cell concentration (SI Figure 1 a,c,d): therefore, for each mouse model, the motile fraction estimate in all caecal samples is the number of motile bacteria per minute of acquisition adjusted by a ratio $C_{feces}/C_{caecum}$ (when both are available) or 1,2.10$^6$/$C_{caecum}$ (when only caecal samples were available). All correction factors and average MFE are provided in SI Table 2.

**SI Methods: detailed information regarding mouse models.**

Samples in this manuscript were all sourced from mouse experiments where they would have otherwise been discarded, without incurring changes in approved protocols. Animals were all bred locally at University Hospital RWTH Aachen. They were housed in individually ventilated cages under specific pathogen-free conditions with 12 h light/dark cycles and water and food (Normal Chow Diet, NCD, by default) available ad libitum. All animal experiments were conducted in accordance with institutional and national guidelines for the care and use of laboratory animals and were approved by the state licensing authorities for animal experimentation, the State Office for Consumer Protection and Nutrition of North Rhine-Westphalia (LAVE, i.e. Landesamt für Verbraucherschutz und Ernährung, North Rhine-Westphalia, Germany). Authorisation numbers are listed in the section of each model if relevant.

**IBD (Inflammatory Bowel Disease) mouse model.** The constitutive interleukin-10 knockout (IL-10 KO) mouse line[1] is widely used for studying chronic colitis due to its heightened susceptibility to inflammatory stimuli and increased incidence of colorectal carcinoma compared to wild-type (WT) controls[1,2]. A total of 16 mice were used, including IL-10 KO and WT animals (n = 8 per genotype; 4 females and 4 males in each group). Animals were co-housed by pairs of same sex. Stool samples that would otherwise have been discarded were collected at multiple time points throughout the study, always by pair (cohoused WT and IL-10 knockout). All samples were obtained prior to sacrifice (10-12 weeks). LAVE animal experiment license n° 81-02.04.2022.A030.

**Salmonella infection mouse model.** 9-week-old female C57BL/6J wild-type mice were infected with wild-type (wt) *Salmonella enterica* subsp. *enterica* serovar Typhimurium (ATCC14028, *S*. Typhimurium). All the mice were pretreated with streptomycin (20 mg, administered by gavage). 24 hours later, one group of mice were orally infected with $10^7$ CFU wt *S.* Typhimurium in PBS, as previously described[3,4]. One group were treated with sterile PBS, used as control mice. Caecum from infected and control mice, that would otherwise have been discarded, were collected at day 2 post infection. LAVE animal experiment license n°84–02.04.2021.A043.

**PSC-IBD (primary sclerosing cholangitis with DSS-induced IBD) mouse model.** *Multidrug resistance protein 2* (*Mdr2*)-deficient mice (*Abcb4*$^{tm1Bor}$)[5] are a well-established model for primary sclerosing cholangitis[6]. They lack the ability to secrete phospholipids from the liver into the bile. This causes increased concentrations of non-micellar components of free bile acids, which act as detergents on the endothelial cells, damaging them and resulting in a leakage of bile into the portal tract. Ultimately, this leads to the activation of myofibroblasts and fibrotic processes in the areas around the inflamed bile ducts. Oral administration of DSS in the drinking water causes IBD with morphological changes resembling human ulcerative colitis[7]. Both sexes were used for the experiment. At 10 weeks of age, acute colitis was induced by administering drinking water enriched with 1.5% DSS (MP Biomedicals, Fountain Parkway, Solon, USA) for a duration of 7 days followed by a subsequent 3-day period of pure drinking water without DSS. For this article, faecal samples that would otherwise have been discarded were collected 3 days before, in the hours following the DSS introduction, after 4 days and 10 days. LAVE animal experiment license n° 81-02.04.2021.A180 and 81-02.04-2023.A162.

**Intestinal neoplasia mouse model.** The mouse model of tamoxifen-induced stabilization of β-catenin in intestinal epithelial cells (*Catnb+/lox(ex3)* B6.Cg-Tg(Vil1-cre/ERT2)23Syr/J, shortened to *Catnb+/lox(ex3)* Vil Cre-ERT)[8,9] is used to study changes in early stages of tumorigenesis. Tamoxifen injection results in continued proliferation of intestinal epithelial cells. This model recapitulates early molecular pathways in colorectal cancer development that affect the entire gut and that we refer to as gut neoplasia. A total of 6 mice (1 male, 5 females) were used, all littermates. Females were cohoused from weaning and throughout the experiment. The mice with Cre+ genotype (n=3) were intraperitoneally injected with 1.5 mg tamoxifen in a total volume of 150µl medicated oil (25 G needle) per mouse to induce neoplastic changes in the gut, which are detectable by day 3 after tamoxifen injection. Control Cre ERT negative mice did not receive tamoxifen. The faecal pellets used in this article were collected on day 13 from all 6 mice, a day before their sacrifice. LAVE animal experiment license n°2024-474_3.

**High-fat diet mouse model.** For induction of obesity and metabolic dysfunction associated with steatotic liver disease[10], B6 wild-type mice or DEREG mice in SPF conditions were placed on a choline-deficient high-fat diet (HFD; cat. no. D05010402, Research Diets) at the age of 10 weeks for a period of 16 weeks. Controls received a normal chow diet (NCD). Faecal samples and caecal samples that would otherwise have been discarded were collected for this article at age 26 week after sacrifice. Both male and female mice were used and female and male data was pooled. LAVE animal experiment license n° M2025-728.

**Flagellin detection deficiency mouse model.** The mouse model with *Tlr5*$^{-/-}$ lacks the gene encoding Toll-like receptor 5, a pattern recognition receptor that recognizes bacterial flagellin and initiates innate immune signalling. Heterozygous littermates (*Tlr5*$^{+/-}$) were used as the control group to minimize the effects of genetic background while retaining comparable breeding and housing conditions.

*Tlr5*$^{+/-}$ and *Tlr5*$^{-/-}$ littermate mice[11] (B6.129S1-*Tlr5tm1Flv*/J, JAX stock #008377) were co-housed from birth and maintained on a normal chow diet. Faecal samples and caecal samples that would otherwise have been discarded were collected for this article after sacrifice at age 8 week by cervical dislocation.

**Motility in gut samples along the digestive tract of healthy mice.** 3 healthy B6 wild-type SPF mice on a normal chow diet (NCD) were sacrificed, the digestive tubes were collected and imported in the anaerobic chamber, then gut content was collected at different anatomical locations. LAVE animal experiment license n°40152A4/50206A4.

**Figures**

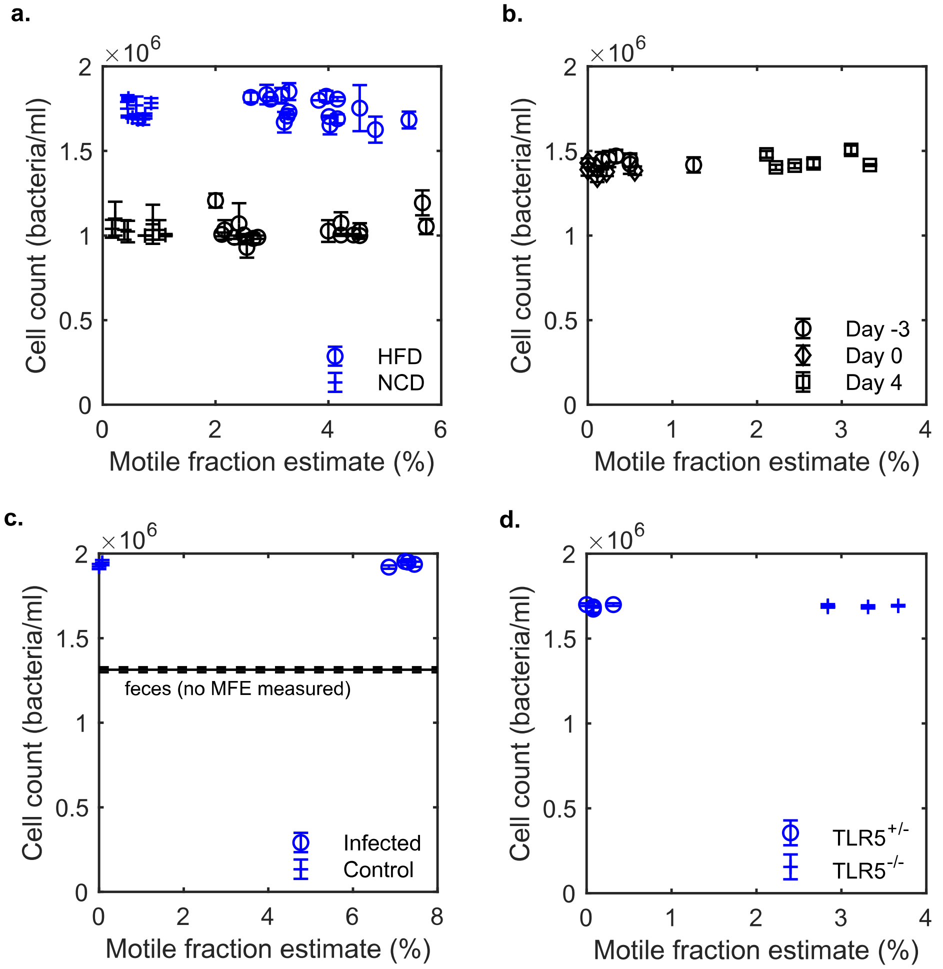


Figure S1 - **Final bacterial concentrations in gut content samples are not a confounding factor in the observed increase of motile fraction estimates.** Plotted are the relationship between motile fraction estimates (MFE, in %) and cell concentrations measured in the samples, for data found in Figure 2 a-f except the IBD model: **(a)** diet model; **(b)** PSC-IBD model across 3 time points relative to introduction of DSS 1.5% in water, **(c)** *Salmonella* Typhimurium infection model and **(d)** flagellin-detection impaired model. Samples from caeca are represented in blue, from faeces in black. Error bars represent standard deviations between replicate cell counting. Each point represents a mouse (at least 3 technical replicates).

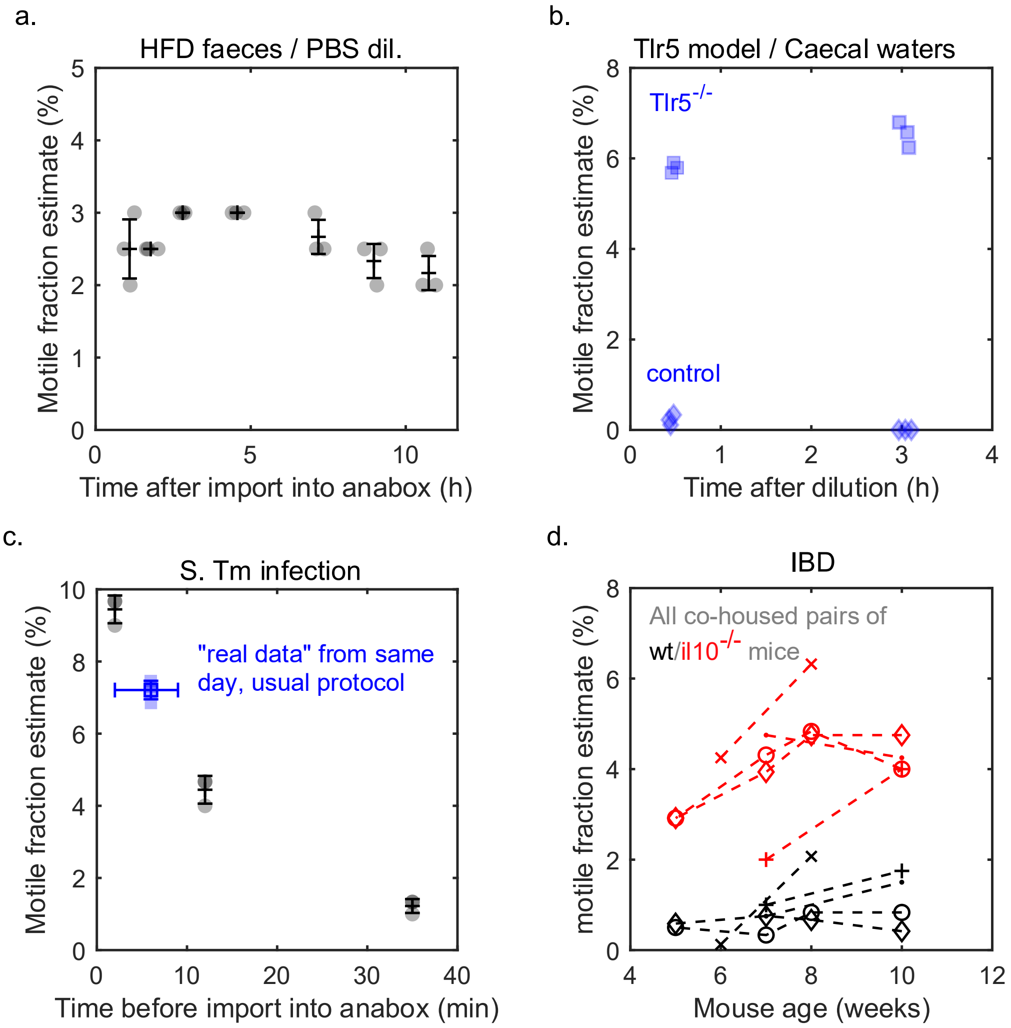


Figure S2 - **Quick import into anaerobic chamber and dilution in PBS ensures stable motile fraction estimates over experimental times, albeit likely underestimating it. (a)** If imported shortly (<10 min) into the anaerobic chamber (here shortened to anabox) and diluted in PBS, the motile fraction estimate of the diluted community (here from 3 faecal pellets of 3 HFD mice) is stable for nearly 11 hours. **(b)** If diluting caecal communities from $Tlr5^{-/-}$ (n=3 mice) and $Tlr5^{+/-}$ mice (n=3 mice) in their respective caecal water (pooled over the 3 mice), we observed changes across 3 hours: an increase in inflamed conditions and a decrease in control conditions (data not shown). Along with the difficulty of preparing fresh caecal/faecal water, this led us to prefer PBS. **(c)** The time spent exposed to air (counted for faeces after dropping, or for caeca after first incision to open the mouse) before import into the anaerobic chamber is critical. Faeces from 3 mice (black) were imported after 2, 12 and 35 minutes then diluted in PBS, displaying a quick loss of motility upon air exposure. Such bias imposed a collection of fresh samples and import within less than 10 minutes in all our experiments (typically under 8 minutes), as well as a parallel collection of controls and inflamed samples to ensure similar treatment. In blue are the results on the same day and same mice as displayed in Figure 2c, following the established protocol. Altogether, our MFE is likely a slight underestimate of the real MFE (i.e. if the import was immediate) due to this incompressible time of collection. **(d)** Further data from the IBD model across mouse ages (5-10 weeks), where cohoused wt (black) and $Il10^{-/-}$ (red, inflamed) mice systematically displayed different MFE across ages. Data in Figure 2a is for 10-week datapoints only.

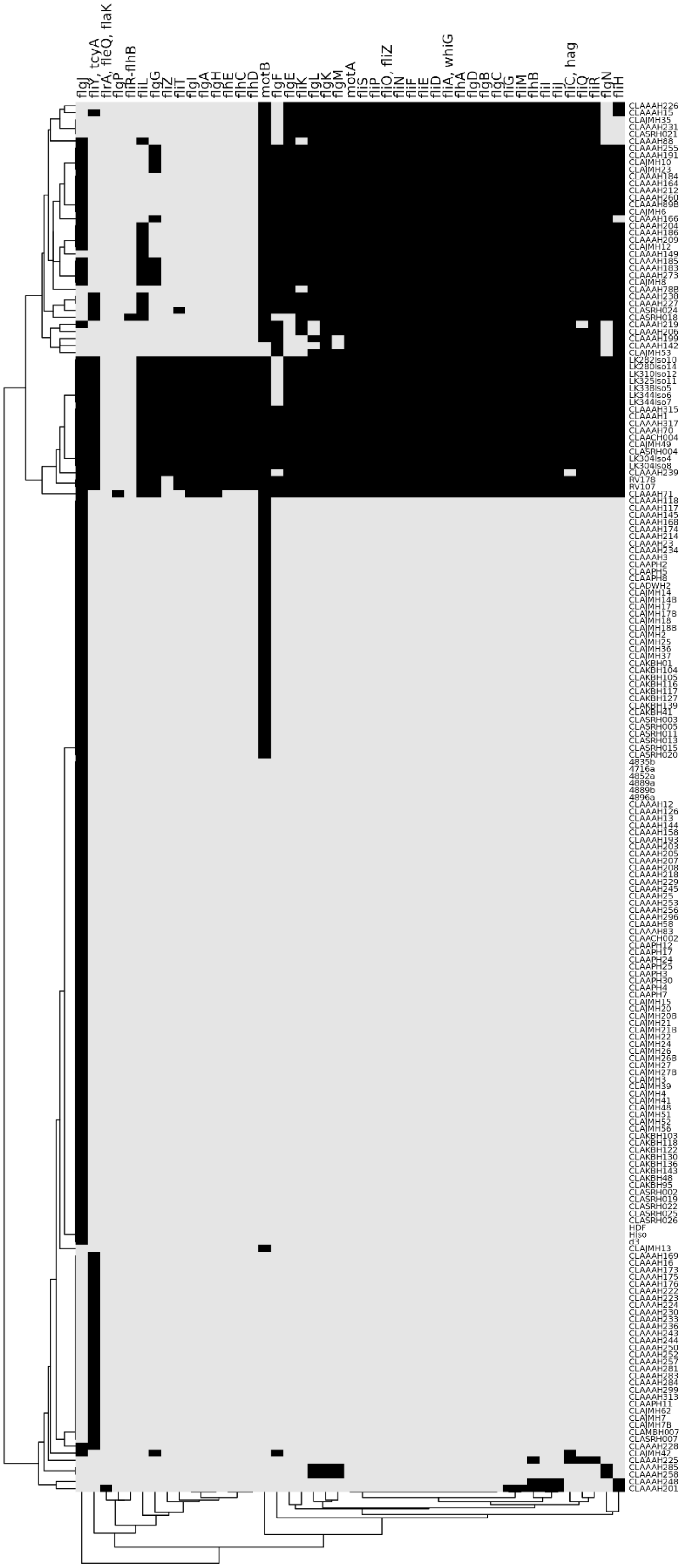

Figure S3 - **Gene screening result with species against the name of flagellar assembly gene(s) detected, if any.** For example, genes flgB,C,D, flhA, motA or fliS systematically associated with motile isolates, while flgJ or motB were not sufficient to predict flagellar motility. Of note, fliC (flagellin) orthologs were found in more than 98% of motile isolates, but also in two isolates that were predicted non-motile and showed no motility (Figure 4d) even upon more extensive experimental testing (SI Figure 4). All 31 isolates from CLAAAH226 (first line) to CLAAAH71included were motile (as validated by SSAPA, Figure 4), the following weren't (as far as we could test). We listed in SI Table 2 the HiBC isolates we used in experiments, the full list of ID of the collection screened here is available at hibc.rwth-aachen.de/ or in Hitch et al., 2025.

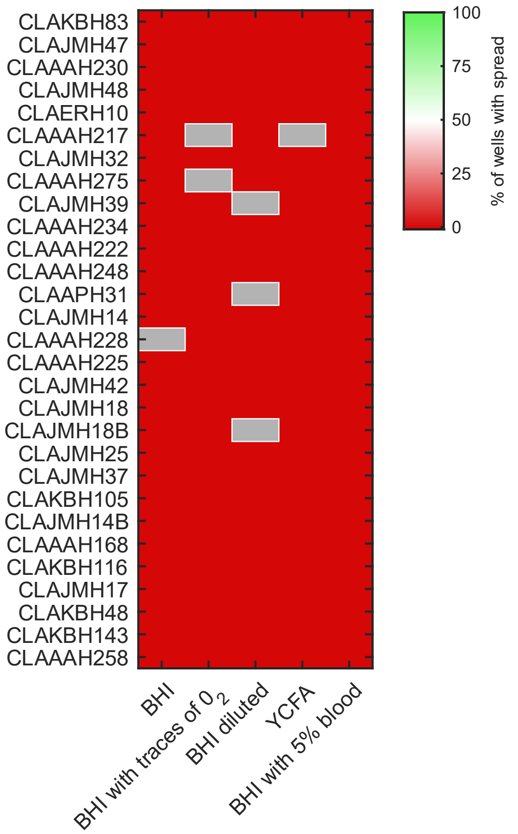


SI Figure 4 - **Further testing the spread in 29 HiBC strains predicted to be non-motile**, over 3 additional growth conditions beyond the BHI, BHI without DTT and YCFA displayed in main text. No spread was observed, which consolidates the hypothesis that these strains are truly non-motile and that the gene screening is a reliable method to predict swimming motility when setting the motility threshold within 8-23 unique genes of the flagellar assembly pathway (KEGG pathway ko02040).

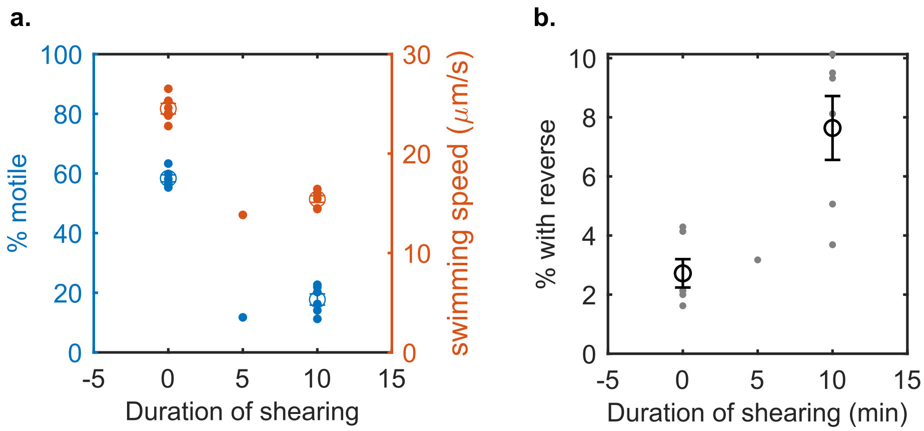


SI Figure 5 - **Shearing of *Salmonella* Typhimurium does not result in more than 10% of the motile population displaying reverse(s) in their trajectory.** *Salmonella* Typhimurium (ATCC14028) was diluted 1:1000 from overnight culture (LB, 30°C, 250 rpm) in 10 ml TB, grown at 30°C until reaching $OD_{600nm}$=1.96, then gently washed (13 min, 1,500 rcf) once and resuspended in 10 ml PBS in a 50 ml Falcon tube. A milk frother (JIMYIU, Amazon ASIN B0D83N2KXC) with a 3-cm whisk fitting inside the tube was used to apply shear to the culture under a biosafety hood. 2 µL of the solution was taken after different time points and diluted in 1ml PBS. Solution were 3D tracked and trajectories with an average speed of at least 10 µm/s were deemed motile. **(a)** The fraction of motile bacteria (blue) and the average swimming speed of motile bacteria (orange) are decreasing, coherent with the expected shear of the peritrichous flagella of *S.* Tm. After 10 minutes of shearing, less than 20 % of the population is motile. **(b)** The reverse-detection analysis does not detect more than 10 % of patterns with at least one reverse, up to 10 minutes of shearing (above which not enough trajectories are available for analysis) where the number of flagella per motile cell is expected to be the lowest. Each point is a technical replicate within the same experiment. Error bars represent the standard error of the mean.

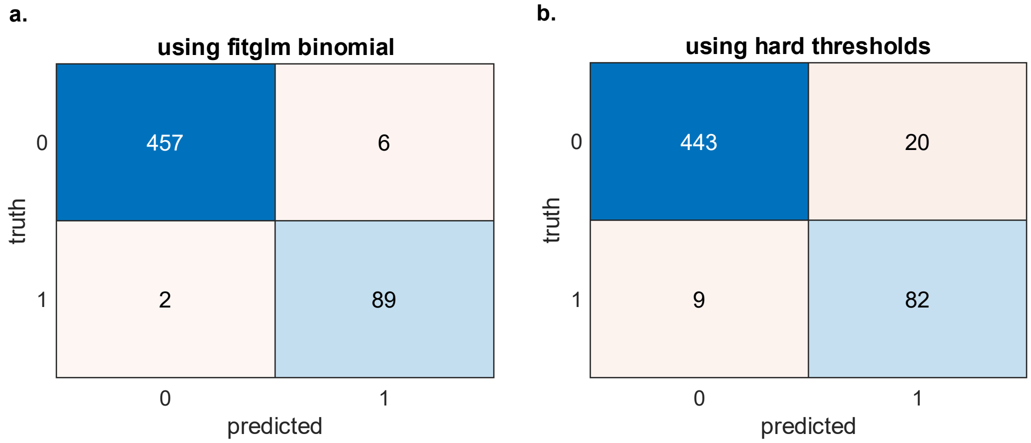


SI Figure 6 - **Model's performances for distinguishing motile (1) from non-motile (0) bacteria tracked in gut contents.** 554 trajectories (463 non-motile, 91 motile) were selected at random from the controls of the diet model **(a)** Confusion matrix of the trained model on 554 trajectories. The binomial generalized linear model (logistic regression) was fitted beforehand to a training dataset of 450 annotated trajectories. **(b)** Confusion matrix of the best combination of hard threshold found for in vitro datasets (median speed, $D_{eff}$, start-end distance, see Methods) on the same 554 trajectories. 0 corresponds to "non-motile", 1 to "motile". The model allowed to reduce false positive from ~4% to 1.4% (a critical point given the large majority of non-motile bacteria in gut contents) and false negative from 9.9% to 2.2%.

**Tables**

**Table S1 - Bacterial species and identifiers, for strains phenotyped and/or used for motile fraction measurements in vitro.**

| Source | Identifier | Species | # of fla. assembly genes |
|---|---|---|---|
| HiBC | CLAAAH255 | *Lachnospira rogosae* | 32 |
| | CLAJMH23 | *Lachnospira rogosae* | 32 |
| | CLAAAH273 | *Waltera acetigignens* | 33 |
| | CLAAAH212 | *Coprococcus hominis* | 31 |
| | CLAAAH78B | *Hominiventricola aquisgranensis* | 29 |
| | CLAAAH15 | *Enterocloster bolteae* | 29 |
| | CLAAAH231 | *Enterocloster citroniae* | 27 |
| | CLAAAH1 | *Escherichia coli* | 42 |
| | CLAAAH239 | *Escherichia coli* | 40 |
| | CLAAAH70 | *Escherichia coli* | 42 |
| | CLAJMH6 | *Lachnospira eligens* | 31 |
| | CLAAAH226 | *Hungatella hathewayi* | 28 |
| | CLAAAH199 | *Intestinimonas aquisgranensis* | 26 |
| | CLAAAH260 | *Lachnospira eligens* | 31 |
| | CLAJMH10 | *Lachnospira hominis* | 32 |
| | CLAAAH191 | *Lachnospira rogosae* | 32 |
| | CLAAAH88 | *Ligilactobacillus ruminis* | 28 |
| | CLAAAH204 | *Roseburia amylophila* | 32 |
| | CLAAAH209 | *Roseburia amylophila* | 32 |
| | CLAAAH238 | *Roseburia intestinalis* | 32 |
| | CLAAAH186 | *Roseburia inulinivorans* | 32 |
| | CLAAAH185 | *Maccoyia intestinihominis* | 33 |
| | CLAJMH12 | *Roseburia amylophila* | 32 |
| | CLAAAH71 | *Selenomonas noxia* | 37 |
| | CLAAAH183 | *Waltera hominis* | 33 |
| | CLAAAH227 | *Robertmurraya yapensis* | 32 |
| | CLAJMH8 | *Waltera intestinalis* | 33 |
| | CLAAAH142 | *Pilosibacter fragilis* | 24 |
| | CLAAAH184 | *Coprococcus hominis* | 31 |
| | RV107 | *Klebsiella aerogenes* | 41 |
| | RV178 | *Klebsiella aerogenes* | 41 |
| | CLAAAH230 | *Bifidobacterium adolescentis* | 1 |
| | CLAJMH48 | *Bacteroides fragilis* | 1 |
| | CLAJMH39 | *Streptococcus thermophilus* | 1 |
| | CLAAAH234 | *Parabacteroides distasonis* | 2 |
| | CLAAAH222 | *Faecalibacterium prausnitzii* | 1 |
| | CLAAAH248 | *Bilophila wadsworthia* | 4 |
| | CLAJMH14 | *Bacteroides thetaiotaomicron* | 2 |
| | CLAAAH228 | *Staphylococcus hominis* | 2 |

| | | | |
|---|---|---|---|
| | CLAAAH225 | *Eisenbergiella tayi* | 4 |
| | CLAJMH42 | *Agathobacter rectalis* | 4 |
| | CLAJMH18 | *Odoribacter splanchnicus* | 2 |
| | CLAJMH18B | *Odoribacter splanchnicus* | 2 |
| | CLAJMH25 | *Bacteroides ovatus* | 2 |
| | CLAJMH37 | *Parabacteroides merdae* | 2 |
| | CLAKBH105 | *Bacteroides faecis* | 2 |
| | CLAJMH14B | *Bacteroides thetaiotaomicron* | 2 |
| | CLAAAH168 | *Bacteroides ovatus* | 2 |
| | CLAKBH116 | *Bacteroides caccae* | 2 |
| | CLAJMH17 | *Parabacteroides distasonis* | 2 |
| | CLAAAH201 | *Desulfovibrio piger* | 7 |
| | CLAKBH143 | *Phocaeicola vulgatus* | 1 |
| | CLAKBH48 | *Phocaeicola vulgatus* | 1 |
| | CLAKBH83 | *Bacteroides fragilis* | 0 |
| | CLAJMH47 | *Bifidobacterium adolescentis* | 0 |
| | CLAERH10 | *Collinsella sp. 2* | 0 |
| | CLAAAH217 | *Blautia fusiformis* | 0 |
| | CLAAAH275 | *Blautia fusiformis* | 0 |
| | CLAJMH32 | *Collinsella sp018373675* | 0 |
| | CLAAPH31 | *Akkermansia muciniphila* | 0 |
| *HFD mouse* | MG14 | *Proteus mirabilis* | - |
| *ATCC* | ATCC12022 | *Shigella flexneri* | - |

**Table S2 - Details of Motility Fraction Estimates (MFEs) in Figure 2a-e and resulting fold increases in inflamed conditions compared to controls.** For each model, the average MFE is displayed along with its standard deviation across mice (each point in Figure 2 corresponding to one mouse, gathering for each mouse at least 3 technical replicate measurements in each type of sample).

| Model | Condition (number of mice) | Sample type | MFE corr. factor | Average motile fraction estimate (MFE±SD, %) | Average fold increase in MFE |
|---|---|---|---|---|---|
| IBD | WT (4) | faeces ● | - | 1.1±0.6 | **3.8** |
| | IL10$^{-/-}$ (4) | | | 4.2±0.3 | |
| Diet | NCD (9) | faeces ● | - | 0.65±0.35 | **5.3** |
| | HFD (17) | | | 3.5±1.3 | |
| | NCD (9) | caeca ■ | 0.6 | 0.60±0.15 | **6.2** |
| | HFD (17) | | | 3.7±0.7 | |
| *Salmonella* Tm Infection | control (6) | caeca ■ | 0.68 | 0.08±0.04 | **102** |
| | infected (8) | | | 8.0±0.9 | |
| Impaired fla. sensing | TLR5$^{+/-}$ (4) | caeca ■ | 0.71 | 0.12±0.14 | **27** |
| | TLR5$^{-/-}$ (4) | | | 3.2±0.4 | |
| PSC-IBD | day -3 before DSS (6) | faeces ● | - | 0.5±0.4 | **5.3** |
| | day +4 after DSS intro. (6) | | | 2.6±0.5 | |
